# ATXN2 polyglutamine expansion impairs QKI-dependent alternative splicing and oligodendrocyte maintenance

**DOI:** 10.1101/2025.08.08.669189

**Authors:** Nesli-Ece Sen, Jaime Eugenin von Bernhardi, Jubril Olamide Adeyemi, Aleksandar Arsović, Júlia Canet-Pons, Antonio J. Miralles, Jana Key, Lorenzo Bina, Vincenzo Romano, Meike Fellenz, Matthias Pietzke, Melanie Halbach, Kay Seidel, Elif Fidan, Zeynep-Ece Kaya-Güleç, Luis E. Almaguer-Mederos, Lindsay A. Becker, Suzana Gispert, Aaron D. Gitler, Thomas Deller, Laurens W. J. Bosman, Chris I. De Zeeuw, David Meierhofer, Leda Dimou, Georg Auburger

## Abstract

**Background:** Polyglutamine (polyQ) tract expansion mutations in Ataxin-2 gene (*ATXN2*) are associated with neurodegenerative diseases spinocerebellar ataxia type 2 (SCA2) and amyotrophic lateral sclerosis (ALS), while the therapeutic reduction of ATXN2 confers strong health-/lifespan extension in models of both disorders. Although the involvement of ATXN2 in peripheral lipid metabolism has been elaborated in *Atxn2* knock-out mice, its impact on nervous system lipid maintenance and a potential influence on oligodendrocytes remains unexplored.

**Methods:** We examine the nervous tissue of an authentic ATXN2 polyQ expansion mouse model in terms of (i) gross morphology of the brain and differential glial affection via immunohistochemical analyses, (ii) spinocerebellar proteome profile via label-free mass spectroscopy and (iii) alternative splicing patterns of oligodendroglial transcripts via quantitative RT-PCR. Finally, electrophysiological recording of sensory response in cerebellar Purkinje cells was performed as a phenotypic measure of demyelination.

**Results:** We demonstrate a massive impairment in myelin maintenance due to ATXN2 polyQ expansion, affecting key oligodendroglial proteins accompanied by their splicing anomalies much earlier than disease manifestation. Oligodendroglial ATXN2 aggregates were documented for the first time in cerebellum, which sequestrated the RNA splicing factor Quaking (QKI). As an outcome of demyelination, our SCA2 model showed a significant delay in response to sensory stimuli.

**Conclusions:** Overall, we provide pioneer evidence of oligodendroglial proteotoxicity leading to myelin maintenance defects in an authentic mouse model of SCA2. Our findings suggest that not only neuronal metabolism, but also that of oligodendroglia depends on ATXN2 and is affected during the disease course. This novel aspect of ATXN2 pathomechanism sheds light on potential outcomes of its therapeutic manipulation, and makes it relevant also for demyelination syndromes next to SCA2 and other polyQ disorders.

## Background

Ataxin-2 (ATXN2) is an evolutionarily conserved, ubiquitously expressed protein involved in translational control and stress response^1,2^. Through its relocalization to stress granules (SG) and its interaction with several RNA-binding proteins (RBPs), ATXN2 modulates transcript quality control versus selective translation^3,4^. Functional studies reveal that ATXN2 contains distinct domains for RNA interaction: the like-Sm (LSm) domain is directly responsible for binding U-rich sequences in the 3′ untranslated regions (3′-UTRs) of target mRNAs and protects against self-aggregation of the protein, while the LSm-associated domain (LSmAD) supports folding and sub-cellular localization including Golgi-associated trafficking^5,6^, and the PAM2 motif enables interaction with poly(A)-binding protein and its associated mRNAs. Large expansions of the N-terminal polyglutamine (polyQ) domain in ATXN2 beyond 34Q are the monogenic cause of widespread neurodegeneration in spinocerebellar ataxia type 2 (SCA2)^7^, whereas intermediate-length expansions of 26-33Q increase the risk for other more common multifactorial neurodegenerative processes. This modifier effect was demonstrated in amyotrophic lateral sclerosis (ALS) and frontotemporal dementia (commonly due to TDP-43 aggregates) where motor neurons are affected, as well as in Parkinson-plus syndromes (usually due to MAP-Tau aggregates) where the basal ganglia are disturbed^8–10^.

A progressively toxic aggregation of ATXN2 underlies the pathology triggered by its polyQ expansion^11^. Notably, N-terminal proteolysis of ATXN2 has been demonstrated to release a polyQ-containing fragment essential to its pathogenicity. This cleavage requires a 17-amino acid motif directly C-terminal to the polyQ stretch, a sequence that is both necessary and sufficient to drive this post-translational modification. The production of this fragment may represent a mechanistic parallel to other polyQ-related diseases such as Huntington’s disease, and adds an additional layer to ATXN2 toxicity, especially relevant for understanding aggregate formation in SCA2^12^. Conversely, silencing of ATXN2 by antisense oligonucleotides or genetic knock-out recently showed immense therapeutic benefit for otherwise fatal ALS and SCA2, extending the lifespan of TDP-43 overexpressing mice up to 10-fold^13,14^. This protective effect appears strongly conserved across phyla, since the silencing of ATXN2 fly ortholog Atx2 ceases neurodegeneration due to tauopathy^10^, and the deletion of ATXN2 yeast ortholog Pbp1 rescues poly(A)-binding-protein (yeast Pab1, mammalian PABP) deletion, yet another SG-associated protein^15^.

Beyond its role in SG dynamics, recent work has demonstrated that ATXN2 is crucial for fast recycling of NOTCH receptors from the ER-Golgi intermediate compartment (ERGIC) to the plasma membrane, promoting rapid activation of Notch signaling. Proteomic mapping revealed that ATXN2 interacts directly with NOTCH1, and its depletion impairs Notch receptor recycling and signaling activation^16^. This agrees with an earlier report documenting the involvement of ATXN2 in epidermal growth factor receptor recycling^17^. ATXN2 thus emerges as an important factor in controlling membrane receptor dynamics, further highlighting its broad impact on cellular processes beyond RNA metabolism.

In order to study ATXN2 pathology with regard to its spatio-temporal evolution and the accompanying molecular dysregulations not only in the nervous system but throughout the body, we recently generated the *Atxn2*-CAG100-knock-in (KIN) mouse model^18^. It was shown by this model that the unstable expansion of the polyQ domain initially impairs the expression and translation of ATXN2, resulting in a partial loss-of-function effect in most peripheral tissues, followed by impaired degradation of expanded ATXN2 leading to its accumulation and aggregation in the nervous system. Mirrored in patients, this process first affects two glutamatergic projections (motor neurons to the spinal cord, and granular neurons in the cerebellum), initially impairing stance, balance, speech articulation, swallowing, gaze and sleep^19^. Despite this preferential onset, the neuropil shrinkage is already massive in all brain regions when the locomotor deficits become detectable. Demyelination appears as early as axonal damage, and eventually widespread neurodegeneration invades the brain. So, SCA2 patients at the terminal stage cannot be easily distinguished on clinical grounds from patients with widespread demyelination^20,21^. It is also important to note that very large expansions or homozygous SCA2 inheritance from both parents will already trigger clinical deficits in childhood, affecting spinocerebellar pathways at the same time as vision, mental development and autonomous nerves. Such severe SCA2 variants present with nystagmus, seizures and muscular hypotonia, exhibiting demyelinating pontocerebellar atrophy or leukoencephalopathy upon brain imaging^22–26^. These findings raise the question if ATXN2 expansions trigger a cerebellum-led multi-system-atrophy (MSA) process with a cell-autonomous affection of oligodendroglia due to cytoplasmic inclusion bodies similar to MSA-C (with SNCA and TAU aggregates) and other Parkinson-plus syndromes^27,28^. Importantly, the neuropathological analyses of SCA2 patient brains showed the severity of glial cytoplasmic inclusions to best predict neurodegeneration, while neuronal nuclear inclusions showed an inverse correlation with neurodegeneration and may therefore play a protective role^29^.

Although both SCA2 and ALS are seen as disorders of selective neurodegeneration, a number of recent findings demonstrated the importance of nutrient metabolism in disease onset and progression^30,31^. Cohort studies showed an earlier disease onset in patients with lower body mass index (BMI), and a significant correlation between BMI reduction rate and disease severity^32,33^. Indeed, the onset and progression of the locomotor deficits in SCA2 mouse models also coincided with accelerated weight loss due to the depletion of peripheral and nervous system lipid stores, *i.e.* cholesterol and N-acetylaspartate (NAA)^18,34–36^. In contrast, obesity with hyperlipidemia and insulin resistance are triggered by the absence of ATXN2 function^37^. Thus, it remains to be determined how the lipid reserves in myelin are depleted by ATXN2 polyQ expansion mechanistically.

In this study, we aimed to deepen these insights by a systematic documentation of ATXN2 polyQ expansion impact on oligodendroglia in order to explore the relevance of glial cells for therapy. Histology showed strong and progressive myelin deficit, suggesting the interference of mutant ATXN2 in oligodendrocyte and myelin maintenance, rather than initial lineage commitment. In addition, global proteome profiles of *Atxn2*-CAG100-KIN cerebellum and spinal cord showed myelin compartment deficits to exceed markers of neuronal atrophy. Further molecular analyses led to the identification of insidious *Mag* and *Plp1* splicing defects, as well as the sequestration of their splicing factor QKI into pathologic aggregates in oligodendrocytes as an upstream event explaining the demyelination phenotype. Electrophysiological recordings in our SCA2 mouse model during whisker pad stimulation showed delayed complex spike responses in cerebellar Purkinje cells presumably due to lower axonal conduction speed. Altogether, our data showed progressive ATXN2 pathology in oligodendrocytes for the first time, which underlies impeded oligodendrocyte maturation and eventual axon-myelin disconnection. We believe, this novel aspect of ATXN2 pathology will encourage the development of diverse therapeutic strategies targeting glial cells in SCA2 and other ATXN2-associated disorders.

## Materials and Methods

### Animals

All animals were kept in individually ventilated cages under a 12-hour light-dark cycle at the Central Animal Facility (ZFE) of the Goethe University Medical School, Frankfurt am Main, Germany. Animals were fed *ad libitum* and were routinely monitored for health. The generation and genotyping of *Atxn2*-CAG100-KIN^18^, *Atxn2*-KO^37^ and ATXN2-Q58-Tg^38^ lines were reported previously. Depletion or transgenic over-expression of the target genes in *Atxn2*-KO and ATXN2-Q58-Tg lines were confirmed by qRT-PCR analyses. Heterozygous mating was performed for the propagation of the colonies. After weaning, age- and sex-matched WT-KIN pairs were housed and aged together, and were further analyzed in parallel employing both sexes. Homozygous KIN animals were used exclusively. All experiments except those involving *in vivo* electrophysiology were approved by the Regierungspräsidium Darmstadt (V54-19c20/15-FK/1083) in accordance with the German Animal Welfare Act, the Council Directive of 24th November 1986 (86/609/EWG) with Annex II and the ETS123 (European Convention for the Protection of Vertebrate Animals). The *in vivo* electrophysiological experiments were approved in advance by the Centrale Commissie Dierproeven (The Hague, The Netherlands, project license no. AVD101002015273) after obtaining positive advice from an independent ethical committee (DEC Consult, Soest, The Netherlands) and overseen by the institutional animal welfare board of Erasmus MC in accordance with the Dutch law on animal experimentation, EU directive 2010/63/EU and institutional guidelines.

### Brain area measurement

Whole brain images of 7 WT (1m/6f) and 8 KIN (4m/4f) animals were taken with Leica stereoscope and areas were quantified with Fiji. After immunolabeling, tile-scan images of whole slices were taken through Keyence fluorescent microscope BZ-9000 using a 10x objective and width was measured manually in Fiji (v1.53c).

### BrdU treatment, perfusion and immunohistochemistry

5-bromo-2′-deoxyuridine (BrdU) was administered orally in drinking water at a concentration of 1 mg/ml with 1% sucrose for 2 weeks. One group of mice (4 WT 2m/2f, 4 KIN 1m/3f) was sacrificed immediately after BrdU treatment, and the other group (8 WT 1m/7f, 9 KIN 5m/4f) was sacrificed at the end of a 4-week wash-off period with regular drinking water. Immunohistochemistry was performed as described earlier^39^. In short: Mice were anesthetized with an intraperitoneal injection of ketamine (100 mg/kg) and xylazine (10 mg/kg) diluted in a 0.9% NaCl solution. Intracardial perfusion was performed with 25 ml PBS and 50 ml 4% PFA/PBS. Brains were isolated and postfixed in 4% PFA/PBS for 2 hours before o/n incubation in 30% saccharose/PBS at 4°C. 30 μm thick free-floating sections were kept in PBS at 4°C. Sections were permeabilized and blocked in 10% goat serum/0.5% Triton-X/PBS for 1 h. Following PBS washes, sections were incubated in primary antibody solutions prepared in blocking buffer o/n at 4°C. Antibodies utilized on BrdU treated samples were: ASPA (Biozol #GTX113389, 1:200), ATXN2 (BD Bioscience #611378, 1:100), GFAP (DAKO #Z0335, 1:500), IBA1 (Synaptic Systems #234004, 1:500), Ki67 (Invitrogen #14-5698-82, 1:100), MBP (Abcam #ab133620, 1:300), MAG (Merck #MAB1567, 1:300), NG2 (Merck #AB5320, 1:500) and OLIG2 (Millipore #MABN50, 1:300), PDGFRα (Thermo Fisher #12-1401-81, 1:250). Following PBS washes, sections were incubated in secondary antibody solutions prepared in blocking buffer for 2 h at RT. Secondary antibodies used in this experiment were: anti guinea pig AlexaFluor-647 (Invitrogen #A21450, 1:500), anti-mouse Cy3 (Dianova #115-165-166, 1:500), anti-mouse AlexaFluor 647 (Invitrogen #A21242, 1:500), anti-rabbit AlexaFluor 488 (Invitrogen #A121206, 1:500), and anti-rabbit Cy3 (Dianova #711-165-152, 1:500). For BrdU detection, after the initial immunostainings, sections were fixed in 4% PFA/PBS for 10 min and incubated in 2 N HCl for 1 h at RT for antigen retrieval. Sections were washed twice in 0.1 M Na_2_B_4_O_7_ and once in PBS. BrdU immunostaining was performed o/n at 4°C using rat αBrdU antibody (Abcam #ab6326, 1:250) followed by anti-rat AlexaFluor (Invitrogen #A21247, 1:500) secondary antibody. Slices were counterstained with DAPI/PBS (1:1000) and washed two times in PBS. Finally, the slices were mounted on glass slides with Aqua-Poly/Mount (Polysciences) and dried overnight. Imaging was done with a Leica SPE inverted confocal microscope using a 40x objective. For cortex acquisition, images of a whole column of the cortex, containing each cortical layer, were taken and three columns per animal were analyzed. For corpus callosum and cerebellar grey and white matter acquisition, at least three field images of lobules IV/V and VI were taken and analyzed. For both acquisitions at least three slices per animal were quantified. Cell quantification was performed manually through Fiji (v1.53c).

### Perfusion and immunohistochemistry

Animals were anesthetized with an intraperitoneal injection of Ketaset (300 mg/kg) and Domitor (3 mg/kg). Intracardial perfusion was performed with 5 min PBS followed by 5 min 4% PFA/PBS. Brains at terminal stage from 3 WT (1m/2f) and 3 KIN (1m/2f) animals, and at pre-onset stage from 2 WT (1m/1f) and 2 KIN (1m/1f) animals were isolated and postfixed o/n in 4% PFA/PBS at 4°C, then were immersed in 30% sucrose for 4 h. 30 μm thick free-floating sections were kept in cryoprotection solution (30% ethylene glycol, 25% glycerin, 0,01% sodium azide in 0.1 M PBS) at −20°C. Sections were washed in 0.1% Triton-X/PBS, permeabilized in 0.3% Triton-X/PBS for 30 min at RT and blocked in 5% goat serum/0.1% Triton-X/PBS for 1 h at RT. Sections were incubated in primary antibody solutions prepared in blocking buffer o/n at 4°C. Primary antibodies utilized in this study were: ATXN2 (BD Biosciences #611378, 1:100; Proteintech #21776-1-AP, 1:200), CALB1 (Cell Signaling #13176, 1:1000), CC1 (Millipore #OP80, 1:50), CNP (Cell Signaling #5664S, 1:100), MAG (Cell Signaling #9043S, 1:100), OLIG2 (Millipore #MABN50, 1:300), PABP (Abcam #ab21060, 1:200), PDGFRα (Novus, #AF1062, 1:100), QKI6 (Neuromab #75-190, 1:200). Following PBS washes, sections were incubated in secondary antibody solutions prepared in blocking buffer for 2 h at RT. Secondary antibodies used in this study were: goat anti-mouse-AlexaFluor-488 (Molecular Probes #A11029, 1:1000), goat anti-mouse-AlexaFluor-546 (Molecular Probes #A11030, 1:1000), goat anti-rabbit-AlexaFluor-488 (Molecular Probes #A11034, 1:1000), goat anti-rabbit-AlexaFluor-546 (Molecular Probes #A11035, 1:1000) together with 1 µg/mL DAPI (Thermo Scientific). After PBS washes, sections were mounted with Lab Vision PermaFluor fluorescent mounting medium (Thermo Scientific) and dried overnight. Imaging was performed with Nikon Eclipse TE2000-E (Nikon) inverted confocal microscope with a 40× objective, and image processing was done with Fiji BioVoxxel software.

### Transmission electron microscopy and image analysis

Four WT (2m/2f) and four KIN (2m/2f) animals were deeply anesthetized with an i.p. overdose of Ketamine (180 mg/kg) and Xylazin (10 mg/kg) and perfused transcardially with 0.9% sodium chloride (NaCl), followed by 4% paraformaldehyde (PFA) / 2.5% glutaraldehyde (Polysciences) in 0.1 M cacodylate buffer (pH 7.4). The cerebella were post-fixed at 4°C o/n in the same fixative. Serial 50 μm-thick sagittal brain sections were cut with a vibratome (Leica VT1000S) and collected in Tris-buffered saline (TBS). Selected sections were washed in 0.1 M cacodylate buffer, osmicated (1% OsO_4_ in cacodylate buffer) for 60 minutes, and dehydrated through an ascending ethanol series (30%, 50%, 1% uranyl acetate [Serva], 70%, 80%, 90%, 100%, 100%), incubated in 100% propylene oxide (2 x 5 minutes), and finally embedded in Durcupan (Sigma-Aldrich). Serial thin sections were cut and collected on single-slot Formvar-coated copper grids and examined with a Zeiss electron microscope (Zeiss EM 900). MyelTracer software^40^ was used for axon diameter and g-ratio measurements with search parameters adjusted to min. size = 50, and max. size = 800. Two images per animal were quantified.

### Post-mortem human samples and Luxol staining

Human cerebellar sections were treated with the modified Heidenhain protocol as described before^41^. An SCA2 patient (biobank ID: GR 245-07, male, ATXN2 CAG repeats: 39/20, disease onset: 32, ex: 51 years) was analyzed together with an age-/sex-matched healthy control.

### Cell culture

Oli-neu cells were cultured at 37°C with 5% CO_2_ in a growth medium consisting of Neurobasal Plus (Gibco), 1X B27 Plus (Gibco), 1% Horse serum (Gibco), 1X ITS liquid media supplement (Sigma), 0.5 µM L-Thyroxine (Sigma) and 50 µg/ml Gentamycin (Sigma) on poly-D-lysine (Gibco) covered flasks or plates. Differentiation was achieved by adding 1 mM dbcAMP (Sigma) into the growth medium for 7 days. Cells were passaged once they reached ∼70% confluence. Mycoplasma contamination was tested regularly.

### Immunocytochemistry

Oli-neu cells were cultured and differentiated as described above on poly-D-lysine (Gibco) coated 12 mm glass cover slips. Following a 30 min oxidative stress induction with 0.5 mM sodium arsenite (NaAsO_2_, Sigma) supplemented in the existing differentiation medium, cells were washed and fixed with 4% PFA/PBS at RT for 20 min. After PBS washes, permeabilization was performed using 0.1% Triton-X-100/PBS for 15 min at RT. Blocking was performed using 3% BSA/PBS solution for 1 h at RT. Primary antibody incubation was done in blocking buffer using ATXN2 (BD Biosciences #611378, 1:100), PABP1 (Abcam #ab21060, 1:300), QKI5 (Merck Millipore #MABN661, 1:100) QKI6 (Neuromab #75-190, 1:200), QKI7 (Neuromab #75-200, 1:100) antibodies o/n at 4°C. Secondary antibody incubation was done in blocking buffer using goat anti-mouse-AlexaFluor-488 (Molecular Probes #A11029, 1:1000), goat anti-mouse-AlexaFluor-546 (Molecular Probes #A11030, 1:1000), goat anti-rabbit-AlexaFluor-488 (Molecular Probes #A11034, 1:1000), goat anti-rabbit-AlexaFluor-546 (Molecular Probes #A11035, 1:1000) antibodies together with 1 µg/mL DAPI (Thermo Scientific) for 1 h at RT in dark. Coverslips were mounted on glass slides with Lab Vision PermaFluor fluorescent mounting medium (Thermo Scientific) and dried overnight. Imaging was performed using Zeiss LSM 510 using a Plan Apochromat 100× objective microscope, and Fiji BioVoxxel software was used for image processing. Plot profiles of SGs were obtained from straightened segments using Straighten tool for each channel separately. Individual signal intensity values per pixel were normalized to average intensity and plotted as normalized intensity (Y) / pixel (X).

### RNA extraction and expression analyses

RNA extraction from ∼20 ug cerebellar and spinal cord tissues, and Oli-neu cell pellet from a T25 flask was performed with TRIzol Reagent (Invitrogen) according to the user manual. DNase digestion and cDNA synthesis from 1 µg of total RNA template was performed using SuperScript IV VILO kit with ezDNase (Thermo Fisher) according to manufacturer’s instructions. Following numbers of mice were used from 3 mo Cb: 8 WT (3m/5f) and 8 KIN (4m/5f), 14 mo Cb: 5 WT (2m/3f) and 3 KIN (2m/1f), 14 mo SC: 5 WT (2m/3f) and 5 KIN (2m/3f), 6 mo Cb: 4 WT (2m/2f) and 4 KO (2m/2f), 16 wk Cb: 4 WT (4f) and 4 Tg (4f), 46 wk Cb: 4 WT (4f) and 4 Tg (4f).

Semi quantitative PCR was performed with cDNA from 25 ng total RNA using AmpliTaq DNA polymerase (Thermo Fisher). The primers utilized in this study were: Mag-exon10-13-Fwd: GTCGCCTTTGCCATCCTGATT, Mag-exon10-13-Rev: TCTCAGATCCCAGGCGCTG, Plp1-Dm20-Fwd: GAAAAGCTAATTGAGACCTA, Plp1-Dm20-Rev: GAGCAGGGAAACTAGTGTGG, *Actb*-Fwd: GGAAATCGTGCGTGACATCAAAG, *Actb*-Rev: CATACCCAAGAAGGAAGGCTGG.

The PCR conditions were 94°C for 4 min, followed by 20 cycles of 94°C for 30 s, 55°C for 45 s and 72°C for 60 s, and a final extension of 72°C for 5 min. Products were resolved on 2% agarose gels at 120V for 30-45 mins, and imaged with Herolab EASY Imaging System. Quantification of signal intensities was performed using ImageStudio software.

Quantitative real-time PCR (qRT-PCR) analyses were performed with StepOnePlus Real-Time PCR System (Applied Biosystems) equipment either with TaqMan or SYBR Green system. For the TaqMan amplification, cDNA from 10ng total RNA was used for each PCR reaction with 0.5 µl TaqMan Assay, 5 µl FastStart Universal Probe Master 2x (Rox) Mix and ddH_2_O up to 10 µl of total volume. The TaqMan Assays utilized for this study were: *Actb* (Mm02619580_g1), *Aspa* (Mm00480867_m1), *Atxn2* (Mm01199894_m1), *Calb1* (Mm00486647_m1), *Cnp* (Mm01306641_m1), *Hapln1* (Mm00618325_m1), *Hapln2* (Mm00480745_m1), *Hapln3* (Mm00724203_m1), *Hapln4* (Mm00625974_m1), *Ina* (Mm00840982_m1), *Mag* (Mm00487538_m1), *Mal* (Mm01339780_m1), *Mbp* (Mm01266402_m1), *Mobp* (Mm02745649_m1), *Mog* (Mm00447824_m1), *Nat8l* (Mm01217217_m1), *Nefh* (Mm01191456_m1), *Nefl* (Mm01315666_m1), *Nefm* (Mm00456201_m1), *Nptn* (Mm00485990_m1), *Plp1* (Mm01297210_m1), *Rtn4* (Mm00445861_m1), *Tbp* (Mm00446973_m1), *Tuba4a* (Mm00849767_s1). The PCR conditions were 50°C for 2 min, 95°C for 10 min, followed by 40 cycles of 95°C for 15 s and 60°C for 1 min. For the SYBR Green system, cDNA from 10ng total RNA was used for each PCR reaction with 2.5 pmol/µl forward and reverse primers, 5 µl qPCR Mastermix Plus for SYBR Green I (Eurogentec, Liège, Belgium) and ddH_2_O up to 10 µl of total volume. The primers utilized in this study were: *L-Mag*-Fwd: AATCGGTCCTGTGGGTGCTG, *L-Mag*-Rev: CGCTGCTTCTCACTCTCATAC, *S-Mag*-Fwd: AATCGGTCCTGTGGGTGCTG, *S-Mag*-Rev: GGGGCTCTCAGTGACAATCC, *Plp1*-Fwd: ATCCCGACAAGTTTGTGGGCAT, *Plp1*-Rev: TATACTGGCAGAGGTCTTGCTA, *Dm-20*-Fwd: TGAGCGCAACGTTTGTGGGCAT, *Dm-20*-Rev: TATACTGGCAGAGGTCTTGCTA, *Cnp1*-Fwd: GGAGCCGGGGCTGACATCTC, *Cnp2*-Fwd: CCCGCAAAGGCGGTGACGGC, *Cnp1-2*-Rev: CTGCCCAAGCTCTTCTTCAGG, *Qki5*-Fwd: TCCTTGAGTACCCTATTGAACCC, *Qki5*-Rev: TAGGTTAGTTGCCGGTGGC, *Qki6*-Fwd: TCCTTGAGTACCCTATTGAACCC, *Qki6*-Rev: TTAGCCTTTCGTTGGGAAAGC, *Qki7*-Fwd: TCCTTGAGTACCCTATTGAACCC, *Qki7*-Rev: GGGCTGAAATATCAGGCATGAC, *Actb*-Fwd: GGAAATCGTGCGTGACATCAAAG, *Actb*-Rev: CATACCCAAGAAGGAAGGCTGG. The PCR conditions were 95°C for 10 min, followed by 40 cycles of 95°C for 15 s and 60°C for 1 min, and a melting curve stage of 95°C for 15 s, 60°C for 1 min and 95°C for 15 s. Gene expression data from qRT-PCR was analyzed using 2^−ΔΔCt^ method with *Tbp* or *Actb* as housekeeping genes.

### Protein isolation and quantitative immunoblots

Cerebellar and spinal cord tissue was homogenized with a motor pestle in 5-10x weight/volume amount of RIPA buffer [50 mM Tris-HCl (pH 8.0), 150 mM NaCl, 2 mM EDTA, 1% Igepal CA-630 (Sigma), 0.5% sodium deoxycholate, 0.1% SDS and Complete Protease Inhibitor Cocktail (Roche)]. Following 30 min incubation at 4°C and 10 min centrifugation at 12000 rpm, supernatant that contains the soluble protein fraction was taken into a fresh tube and used in further analyses. Following numbers of mice were used from 3 mo Cb: 4 WT (4f) and 4 KIN (4f), 14 mo Cb: 5 WT (2m/3f) and 5 KIN (2m/3f), 14 mo SC: 5 WT (2m/3f) and 5 KIN (2m/3f). For PN/Urea fractionation, fresh cerebella were processed as described earlier^18^. Three WT (3m) and three KIN (3m) animals were used. Oli-neu cultures were grown, differentiated and starved in T25 flasks coated with PDL (Gibco). Cell pellets from control and starvation samples (6 h in HBSS medium) were homogenized in 100 µl RIPA buffer, incubated 20 min at 4°C and sonicated. 20 µg of total protein from mouse tissues and 10 µg from Oli-neu cells were mixed with 2x loading buffer [250 mM Tris-HCl pH 7.4, 20% Glycerol, 4% SDS, 10% 2-Mercaptoethanol, 0.005% Bromophenol blue], incubated at 90°C for 2 min, separated on polyacrylamide gels and transferred to Nitrocellulose membranes (GE Healthcare). The membranes were blocked in 5% BSA/TBS-T for 1 h at RT, and incubated overnight at 4°C with primary antibodies against ATXN2 (Proteintech #21776-1-AP, 1:500), ACTB (Sigma #A5441, 1:10000), CALB1 (Cell Signaling #13176, 1:2500), CNP (Cell Signaling #5664S, 1:1000), MAG (Cell Signaling #9043S, 1:500), MBP (Merck #05-675, 1:250), MOG (Abcam #ab32760, 1:500), NEFH/M (Proteintech #18934-1-AP, 1:500), NEFL (Cell Signaling #2837S, 1:1000), NPTN (Alomone Labs #ANR-090, 1:500), PLP1 (Abcam #ab28486, 1:1000), TUBA4A (Aviva Systems #ARP40179-P050, 1:500), QKI5 (Merck Millipore, #MABN661, 1:500), QKI6 (Neuromab, #75-190, 1:2500) and QKI7 (Merck Millipore, #AB9908, 1:1000). Fluorescent labeled secondary goat anti-mouse (IRDye 800CW, Li-Cor, 1:10000) and goat anti-rabbit (IRDye 680RD, Li-Cor, 1:10000) antibodies were incubated for 1 h at RT. Membranes were visualized using Li-Cor Odyssey Classic instrument. Quantification of signal intensities was performed using ImageStudio software.

### Proteomics sample preparation with label-free quantification

Six homozygous *Atxn2*-CAG100-KIN and eight WT mice at terminal stage (14 mo) were used for proteome profiling. Cerebellum and spinal cord were dissected and snap frozen in liquid nitrogen. Between 20 and 40 mg of each tissue were homogenized under denaturing conditions with a FastPrep (two times for 60 s, 6.5 m x s^-1^) in 1 mL of a fresh buffer containing 3 M guanidinium chloride (GdmCl), 10 mM tris(2-carboxyethyl)phosphine, 20 mM chloroacetamide and 100 mM Tris-HCl pH 8.5. Lysates were boiled at 95°C for 10 min in a thermal shaker, followed by sonication for 10 min and centrifuged at 16,000 rcf for 10 min at 4°C. The supernatant was transferred into new protein low binding tubes (Eppendorf, Germany). 50 µg protein per sample were diluted to 1 M GdmCl by adding 10% acetonitrile and 25 mM Tris, 8.5 pH, followed by a Lys-C digestion (Roche, Basel, Switzerland; enzyme to protein ratio 1:50, MS-grade) at 37°C for 2 h. This was followed by another dilution to 0.5 M GdmCl and a tryptic digestion (Roche, 1:50) at 37°C, at 800 rpm, and o/n. Subsequently, peptides were desalted with C18 columns and reconstituted in 2% formic acid in water and further separated into five fractions by strong cation exchange chromatography (SCX, 3M Purification, Meriden, CT). Eluates were first dried in a SpeedVac, then dissolved in 5% acetonitrile and 2% formic acid in water, briefly vortexed, and sonicated in a water bath for 30 seconds prior to injection to nano-LC-MS/MS.

### LC-MS/MS instrument settings for shotgun proteome profiling and data analysis

LC-MS/MS was carried out by nanoflow reverse-phase liquid chromatography (Dionex Ultimate 3000, Thermo Scientific) coupled online to a Q-Exactive HF Orbitrap mass spectrometer (Thermo Scientific), as reported previously^42^. Briefly, the LC separation was performed using a PicoFrit analytical column (75 μm ID × 50 cm long, 15 µm Tip ID; New Objectives, Woburn, MA) in-house packed with 3 µm C18 resin (Reprosil-AQ Pur, Dr. Maisch, Ammerbuch, Germany). Peptides were eluted using a gradient from 3.8 to 38% solvent B in solvent A over 120 min at a 266 nL per minute flow rate. Solvent A was 0.1 % formic acid and solvent B was 79.9% acetonitrile, 20% H_2_O, and 0.1% formic acid. For the IP samples, a one-hour gradient was used. Nanoelectrospray was generated by applying 3.5 kV. A cycle of one full Fourier transformation scan mass spectrum (300−1750 m/z, resolution of 60,000 at m/z 200, automatic gain control (AGC) target 1 × 10^6^) was followed by 12 data-dependent MS/MS scans (resolution of 30,000, AGC target 5 × 10^5^) with a normalized collision energy of 25 eV.

Raw MS data were processed with MaxQuant software (v2.1.0.0) and searched against the mouse proteome database UniProtKB with 55,366 entries, released in March 2021. Parameters of MaxQuant database searching were a false discovery rate (FDR) of 0.01 for proteins and peptides, cysteine carbamidomethylation was set as fixed modification, while N-terminal acetylation and methionine oxidation were set as variable modifications. The mass spectrometry data have been deposited to the ProteomeXchange Consortium (http://proteomecentral.proteomexchange.org) via the PRIDE partner repository^43^ with the dataset identifier PXD036124.

Proteomics data were filtered to include only proteins detected in at least three replicates per genotype. Fold-changes and p-values were then calculated for each tissue independently. Commonly identified proteins between cerebellum and spinal cord were obtained by matching the calculated fold-changes in both tissues using the gene names. In a few cases multiple protein groups matched to the same gene name, here the fold-changes were averaged. Among the commonly identified proteins, those with similar dysregulation levels in both tissues by 20% were filtered to study commonly dysregulated subset of the proteome in two affected tissues. Protein-protein interaction network of the commonly downregulated proteins was constructed using STRING database v11.5 (https://string-db.org/). Network parameters were imported into Cytoscape software v3.9.1, onto which the fold change values from the proteome data were integrated. Using stringApp in Cytoscape, pathway enrichment analysis was conducted with the default settings. The largest subnetwork is displayed with node colors indicating the level of downregulation, and surrounding donuts indicating the pathway each protein contributes to. Pathway enrichment analysis results are provided in **Table S1**.

### Surgical procedures and in vivo electrophysiology

Mice of 8-9 months old (5 WT [3m/2f] and 9 KIN [4m/5f]) received a magnetic pedestal for head fixation, attached to the skull above bregma using Optibond adhesive (Kerr Corporation) and a craniotomy to expose cerebellar crus 1 and crus 2 on the right side under isoflurane anesthesia (2%–4% v/v in O_2_). The craniotomy was cleaned and afterward covered with Kwik-Cast (World Precision Instrument, Sarasota, FL, USA). Postsurgical pain was treated with carprofen (Rimadyl, Pfizer), buprenorphine (‘‘Temgesic’’ Indivior, Richmond, VA, USA), bupivacaine (Actavis, Parsipanny-Troy Hills, NJ, USA) and lidocaine (Braun). After 3 days of recovery, mice were habituated to the recording setup during at least two daily sessions of approximately 45 min each. In the recording setup, they were head-fixed using the magnetic pedestal. Electrophysiological recordings were performed in awake mice using quartz-coated platinum/tungsten electrodes (2– 5 MΩ, outer diameter = 80 µm, Thomas Recording, Giessen, Germany) placed in an 8 × 4 matrix (Thomas Recording), with an inter-electrode distance of 305 µm. Prior to the recordings, the mice were lightly anesthetized with isoflurane to remove the dura, bring them in the setup and adjust all manipulators. Recordings started at least 60 min after termination of anesthesia and were made in crus 1 and crus 2 ipsilateral to the side of the whisker pad stimulation at a minimal depth of 500 µm. The electrophysiological signal was digitized at 25 kHz, using a 1–6000 Hz band-pass filter, 22x pre-amplified and stored using a RZ2 multi-channel workstation (Tucker-Davis Technologies, Alachua, FL). Air-puff stimulation to the whisker pad was applied with a frequency of 0.5 Hz at a distance of approximately 3 mm at an angle of approximately 35° (relative to the body axis). The puffs were delivered using a tube with a diameter of approximately 1 mm with a pressure of ∼2 bar and a duration of 30 ms.

### Statistical analyses

Unless stated otherwise and except for global proteome and electrophysiology data, all statistical analyses between WT and mutant mice were performed using unpaired Student’s t-test with Welch’s correction using GraphPad Prism 10 software. Graphs display mean values and standard error of the mean. T 0.05<p<0.1, * p<0.05, ** p<0.01, *** p<0.001, **** p<0.0001.

## Results

### Profound white matter atrophy in cerebellum due to ATXN2 toxicity

The well-characterized disease course of KIN mice starts with frequent falls on Rotarod at 3-4 months of age, followed by a progressive loss of body weight at 6-12 months, before the animals have to be sacrificed when the weight loss per month becomes excessive at around 14 months. We simplify the KIN disease course into pre-onset (<3 months), early symptomatic (3-6 months), late symptomatic (6-12 months) and terminal (12-14 months) stages taking the Rotarod performance and weight profiles into consideration (**Supp. Figure S1a)**^18^. Throughout this study, we used 9-10 month-old animals to assess the dynamic changes in the brain architecture, 14 month-old animals to assess RNA and protein dysregulations at the terminal stage and 3 month-old animals to investigate which of these changes occurred in the pre-onset stage and could be subject to therapeutic manipulations.

Gross morphological assessment of the KIN brain at late-symptomatic stage showed a significant reduction in both cerebellar and cerebral area (20% and 12% respectively, **Figure 1a-b**). In cerebellar gray matter (GM), only the molecular layer area where the glutamatergic parallel fiber axons synapse with Purkinje neuron dendrites was found to be reduced, while the granule neuron layer which is dominated by myriads of nuclei with little cytoplasm showed no shrinkage (**Figure 1c-d**). A stronger reduction was observed in the cerebellar white matter (WM) area, thus leading to a significant decrease in the ratio of WM versus total GM area (**Figure 1e-f**), indicating that demyelination is at least as strong as neuropil loss. In the cerebrum, thickness of cortical GM was significantly reduced at variable distances from the midline (**Figure 1g-i**). Furthermore, thickness of the cortical WM (corpus callosum) showed a similarly strong affection (**Figure 1g, h, j**), leading to no significant change in the ratio of WM versus GM size (**Figure 1k**). Overall, quantitative morphology showed GM and WM to be similarly affected in cerebrum, with a tendency towards a stronger WM affection, *i.e.* demyelination, in cerebellum.

**Fig. 1:**
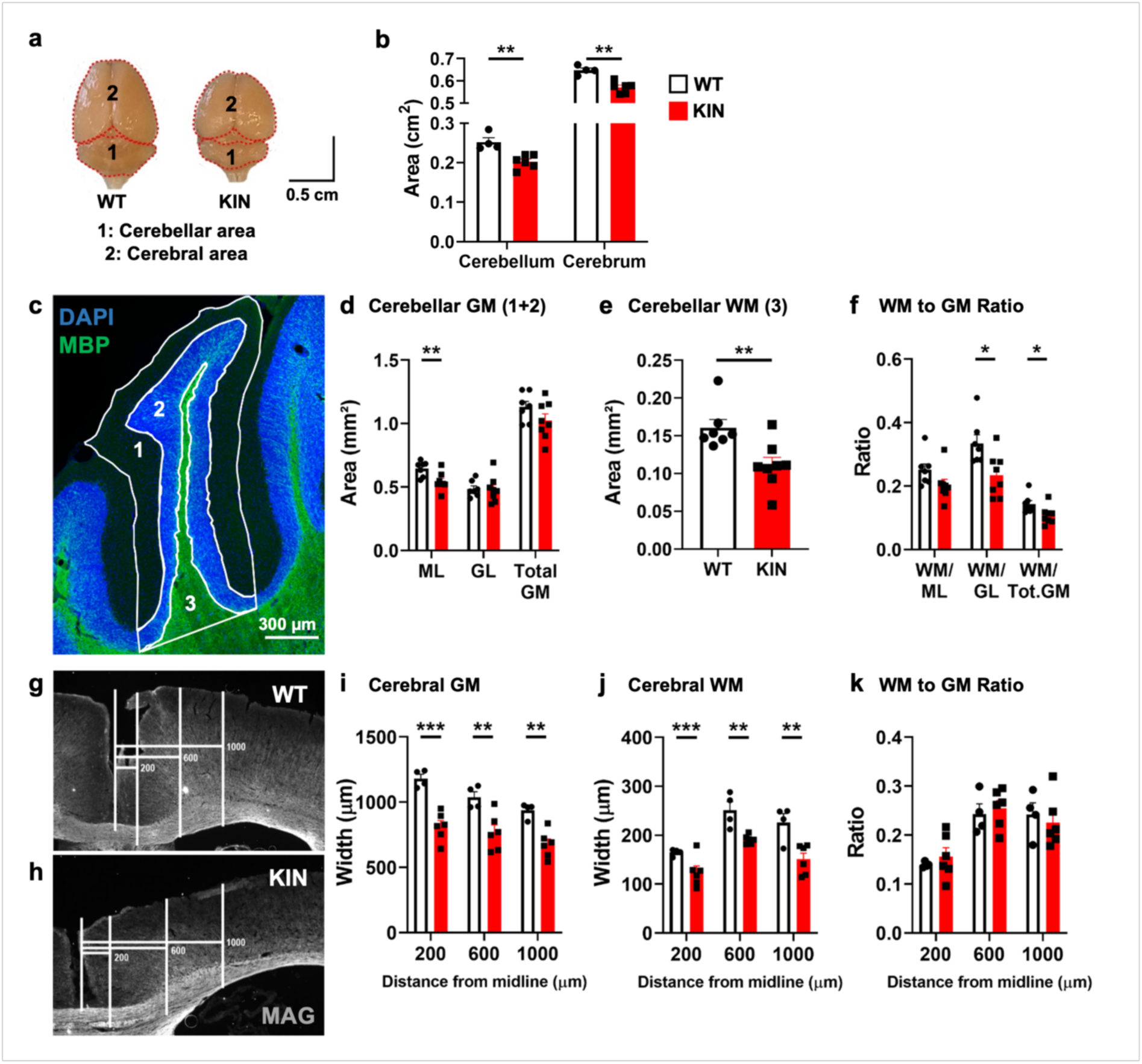
Prominent effect of ubiquitous ATXN2 toxicity on cerebellar white matter. **a,** Representative images of 10-months old WT (left) and KIN mutant (right) brains, showing the macrostructural differences between the two groups. Dotted red lines enclose the compared and quantified areas: cerebellum (1) and cerebrum (2). **b,** Quantification of cerebellar and cerebrum areas shown in panel a. **c,** Representative tile-scan of the simple lobe of the cerebellum and area enclosed by straight lines are the (1) molecular (ML), and (2) granular layers (GL), as well as the (3) white matter arbor vitae (WM), which were measured to identify differences between WT and KIN mice. Slices were immunolabeled against MBP (green) and counterstained with DAPI (blue). **d,e,** Raw measures of the ML, GL, total GM (ML+GL), and WM in WT and KIN mice. **f,** Graph depicting the ratio between the WM and ML, GL or total GM (ML plus GL) in WT and KIN mice. **g,h,** Representative tile-scan images of the cortical GM and WM of the cerebrum. Horizontal straight lines show the distance from the midline (200, 600, and 1000 µm) and vertical straight lines show the width measured. Histological slices were immunolabeled against MAG (white). **i,j,** Quantification of the width of the cortex and corpus callosum at different distances from the midline (200, 600, and 1000 µm). **k,** Graph depicting the ratio between the WM and GM at different distances (200, 600, and 1000 µm). Each data point represents a single animal.

### Demyelination affects axons of various diameters in cerebellum

Transmission electron microscopy (TEM) was employed to confirm that the WM shrinkage caused by ATXN2 toxicity was indeed due to demyelination. Visualization of middle cerebellar peduncles at late-symptomatic stage showed reduced myelin thickness in KIN mice (**Figure 2a, b**). The extent of demyelination in KIN animals was quantified using g-ratio on individual axons, revealing collectively higher g-ratios yielding a significant difference between the slopes of regression lines for WT (r = 0.3735) and KIN (r = 0.5727) samples (**Figure 2c**). Then, we asked whether axons of a certain diameter were selectively affected by ATXN2 toxicity-induced demyelination. Grouping together various axon diameters (<0.6 µm, between 0.6-0.9 µm, between 0.9-1.2 µm, >1.2 µm) also showed significantly higher g-ratios for KIN animals throughout the spectrum (**Figure 2d**). The number of axons in these subgroups, however, did show a difference between WT and KIN animals. We observed a significantly lower percentage of larger axons with >1.2 µm diameter in KIN (38.15%) compared to WT (55.26%) cerebella. While the axons in 0.9-1.2 µm interval did not show a drastic difference (WT: 23.64% / KIN: 25.81%), KIN cerebella had higher percentage of smaller diameter axons at 0.6-0.9 µm (WT: 17.99% / KIN: 29.62%) and < 0.6 µm intervals (WT: 3.10% / KIN: 6.41%), albeit without statistical significance (**Figure 2e**). In addition to the axons in the middle cerebellar peduncle, axons running through the WM in cerebellar lobules were visualized in sagittal sections using immunohistochemical assessment of Calbindin-1 (CALB1), an established marker of cerebellar Purkinje neurons. In accordance, CALB1 staining revealed irregular axon bundles in KIN tissue at terminal disease stage as a known outcome of demyelination (**Figure 2f, g**). Furthermore and more importantly, a similar phenomenon was observed in the *post-mortem* cerebellum of a SCA2 patient (**Figure 2h, i**), where axons showed irregular sprouting suggestive of interrupted myelin sheaths^44^. Together, these data establish the presence of significant demyelination in KIN cerebella already in the late-symptomatic stage, with axons of various diameters affected equally, together with evidence for axonal sprouting at the terminal stage in KIN animals and in *post-mortem* SCA2 patient tissue.

**Fig. 2:**
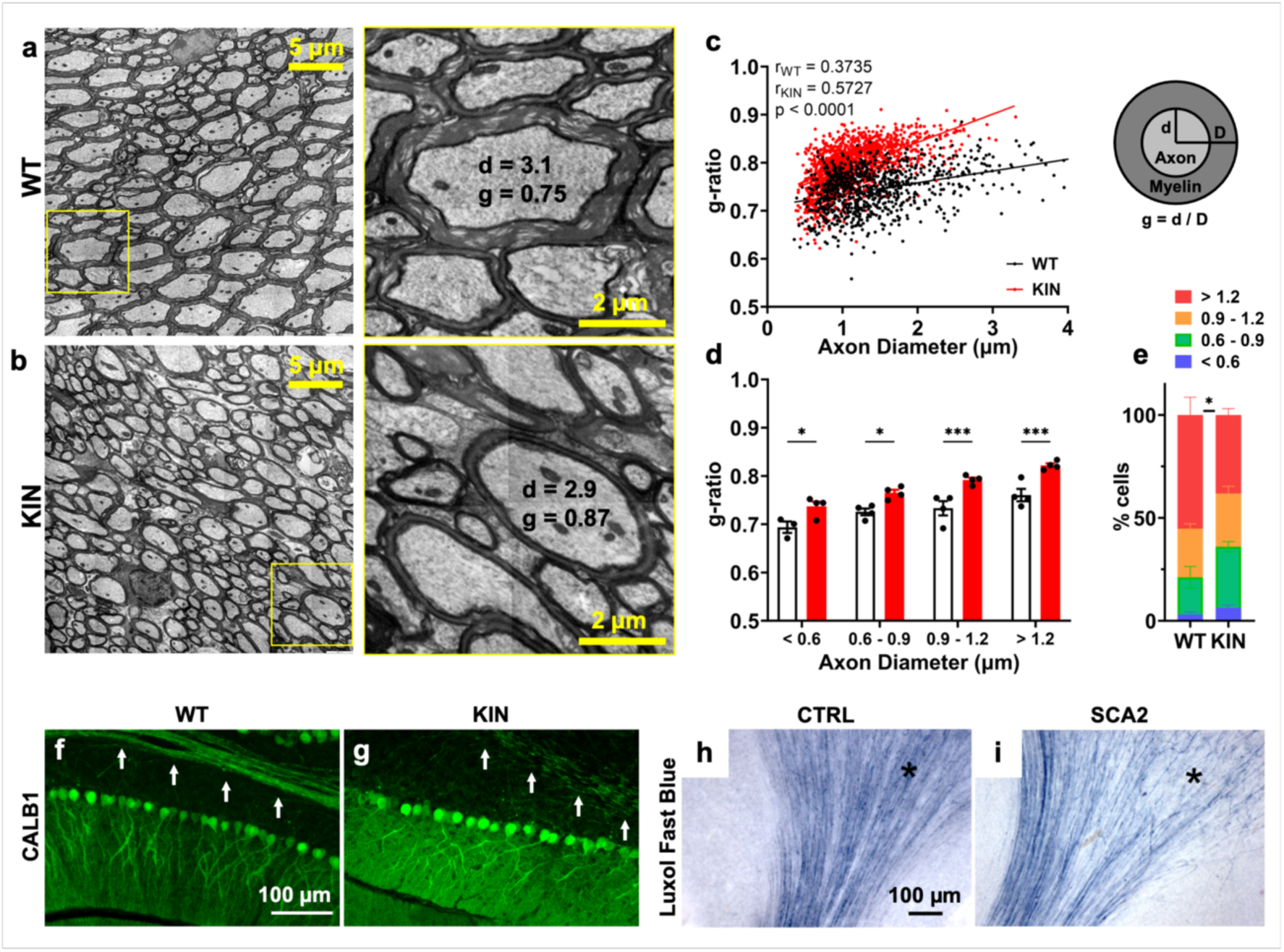
Significant demyelination in KIN cerebellum affects all axon diameters. **a,b,** Representative images of mid-cerebellar peduncle showing pronounced demyelination in late-symptomatic KIN mice (9 mo). **c,** Quantification of g-ratios (explained in the scheme on the right) for each axon was done to assess the rate of demyelination from the TEM images. Collectively higher g-ratios was evident in KIN mice with a significantly different regression (r_WT_ = 0.3735 vs. r_KIN_ = 0.5727, p < 0.0001). Each data point represents a single axon from 4 WT and 4 KIN animals. **d,** Average g-ratios were quantified for axons with various diameters for each animal, showing significantly high g-ratios, therefore demyelination, in KIN mice across all axon diameters. Each data point represents a single animal. **e,** Percentage of cells with various axon diameters were quantified for 4 WT and 4 KIN animals, showing a significant reduction in the number of larger axons in KIN mice. **f,g,** Calbindin-1 (CALB1) immunohistochemistry in WT and KIN cerebella at terminal stage (14 mo) shows Purkinje cell soma to be intact, however sprouting of the axon bundles due to loss of myelin is observed in KIN mouse (white arrows). **h,i,** Axonal sprouting is also observed upon myelin staining with Luxol Fast Blue in *post-mortem* SCA2 patient cerebellum compared to an age- and sex- matched control.

### ATXN2 toxicity impairs oligodendrocyte maintenance and adult oligodendrogenesis in cerebellum

A targeted evaluation of distinct glia types was undertaken in order to examine the differential affection of each cell population, and to assess if compensatory proliferation and glial activation occurs. As NG2-glia can differentiate into mature oligodendrocytes throughout life^45,46^, we wanted to study the generation of oligodendrocytes in the late-symptomatic phase. Therefore, two groups of mice received bromodeoxyuridine (BrdU) in the drinking water over 2 weeks, and in one of the groups this was followed by a 4-week wash-off period to trace the progeny of these proliferating cells (**Supp. Figure S1b**).

In KIN cerebellar WM, significantly elevated microglial activation (**Supp. Figure S2a-e**) and astrogliosis (**Supp. Figure S2f-j**) were observed as expected in both treatment groups. The number of both total and proliferating NG2-glia as oligodendrocyte progenitor cells (OPCs) were also found increased in KIN animals (**Figure 3a-d, i, j**), suggesting an inherent urge to counteract demyelination. Actively proliferating Ki67+/PDGFRα+ oligodendrocyte progenitors also showed elevated numbers in KIN tissue (**Figure 3k**). Tracing the maturation of these newly generated cells (in 2-wk BrdU + 4-wk wash-off group) revealed an immense induction of mature oligodendrocyte production in the KIN cerebellum (**Figure 3e-h, l)**, possibly as a result of compensatory replacement. Despite this strong induction, the number of mature oligodendrocytes was comparable between WT and KIN animals four weeks after labeling (**Figure 3l**). This indicates that the maintenance of newly generated ASPA+ mature oligodendrocytes failed in KIN tissue due to ATXN2 toxicity. When we compared these results to the cerebellar GM, KIN mice also showed higher microglial activation and astrogliosis in both treatment groups (**Supp. Figure S1c-d**). The number of total and proliferating NG2-glia were comparable between WT and KIN animals (**Supp. Figure S1e-f**), however with significantly higher number of mature oligodendrocytes present in KIN mice (**Supp. Figure S1g**). KIN mice also showed a higher number of newly generated oligodendrocytes, yet without statistical significance due to high variation (**Supp. Figure S1h**), again suggesting an increased need for oligodendrocyte replacement, however to a lesser extent due to the lower physiological myelination levels in the GM. These findings were mirrored in cerebrocortical GM and WM, with significantly elevated microgliosis (**Supp. Figure S3a, g**), astrogliosis (**Supp. Figure S3b, h**), and a push towards oligodendrocyte replacement more prominent in the WM (**Supp. Figure S3c-f, i-l**).

**Fig. 3:**
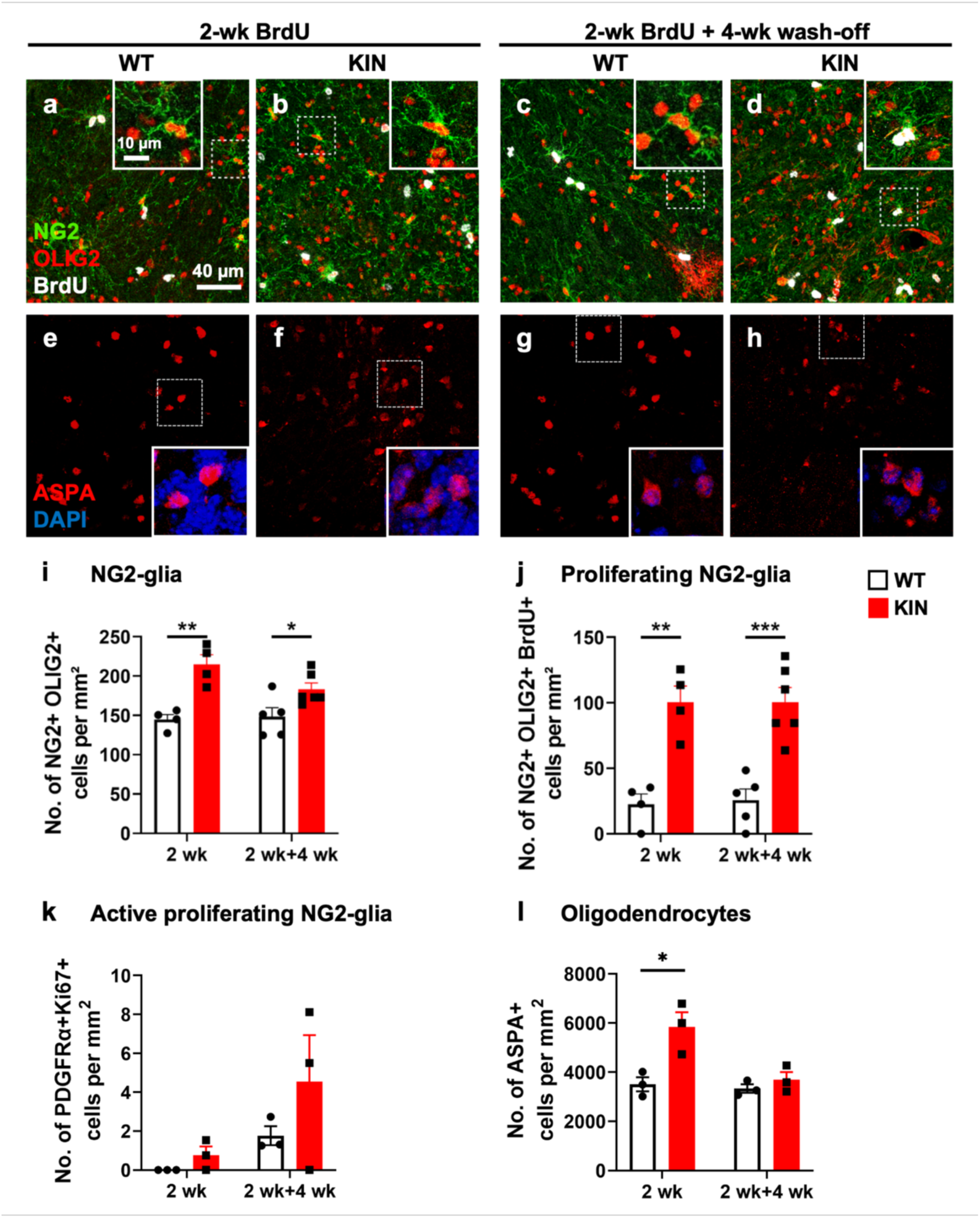
ATXN2 toxicity hinders oligodendrocyte maturation in cerebellar white matter. Immunohistochemical analyses of NG2 and OLIG2-positive cells in cerebellar white matter revealed significantly increased numbers of oligodendrocyte progenitors demonstrated by total **(a-d, i)** and proliferating NG2-glia counts **(a-d, j)** in late-symptomatic stage. The number of PDGFRα and Ki67-positive actively proliferating NG2 glia were also considerably high in KIN cerebella in both treatment groups (**k**), however without statistical significance. Number of mature oligodendrocytes assessed by ASPA-positive cell quantification was found significantly increased in KIN animals immediately after BrdU administration, however their number after the 4-week wash-off period were comparable between WT and KIN animals (**e- h, l**). Each data point represents a single animal.

Jointly, these data indicate an overall inflammatory response to ATXN2 expansion toxicity in terms of microglial activation and astrogliosis throughout the brain. While oligodendrocytic parameters were understandably more subtle in both cerebral and cerebellar GM, profound elevations were observed in oligodendrocyte progenitor numbers, their proliferation, as well as newly generated oligodendrocyte numbers in KIN WM, which fails to persist after four weeks. The fact that WM atrophy and demyelination are evident despite the compensatory efforts in gliogenesis suggests the interference of mutant ATXN2 in maintenance of mature oligodendrocytes or in their full differentiation into myelinating cells, rather than earlier steps involving progenitors and their proliferation.

### ATXN2 is expressed in oligodendrocytes and forms cytosolic aggregates in disease

To address whether the demyelination is a secondary outcome of neuronal/axonal dysregulations, or if it could originate from the ATXN2 pathology within the oligodendrocytes, we assessed the presence of ATXN2 protein and aggregates in oligodendroglia. In differentiated Oli-neu cells, ATXN2 showed diffuse cytosolic localization under normal conditions, and relocalized into PABP-positive stress granules (SGs) under oxidative stress (**Figure 4a**). Furthermore, both transcript and protein abundance of ATXN2 in oligodendrocytes increased upon starvation stress (**Figure 4b, c**), in line with previous reports in other cell types^3,47^. Examination of mouse cerebella also showed diffuse cytosolic localization of WT ATXN2, while the mutant protein was found in cytosolic aggregates in OLIG2+ oligodendroglia (**Figure 4d**). To further examine ATXN2 expression across different stages of the oligodendrocytic lineage, its localization was analyzed in OPCs and mature oligodendrocytes using PDGFRα and CC1 as markers, respectively (**Figure 4e**). The number of ATXN2-expressing OPCs were not changed despite the significantly higher number of total and ATXN2-negative OPCs in KIN mice (**Figure 4f**). In contrast, the number of mature oligodendrocytes expressing ATXN2 was found increased in KIN mice, despite an overall decrease in total and ATXN2-negative mature oligodendrocytes (**Figure 4g**). Quantification of all ATXN2-positive cells in cerebellar WM showed a significant portion to be oligodendrocyte lineage cells in both WT and KIN mice (**Figure 4h**).

**Fig. 4:**
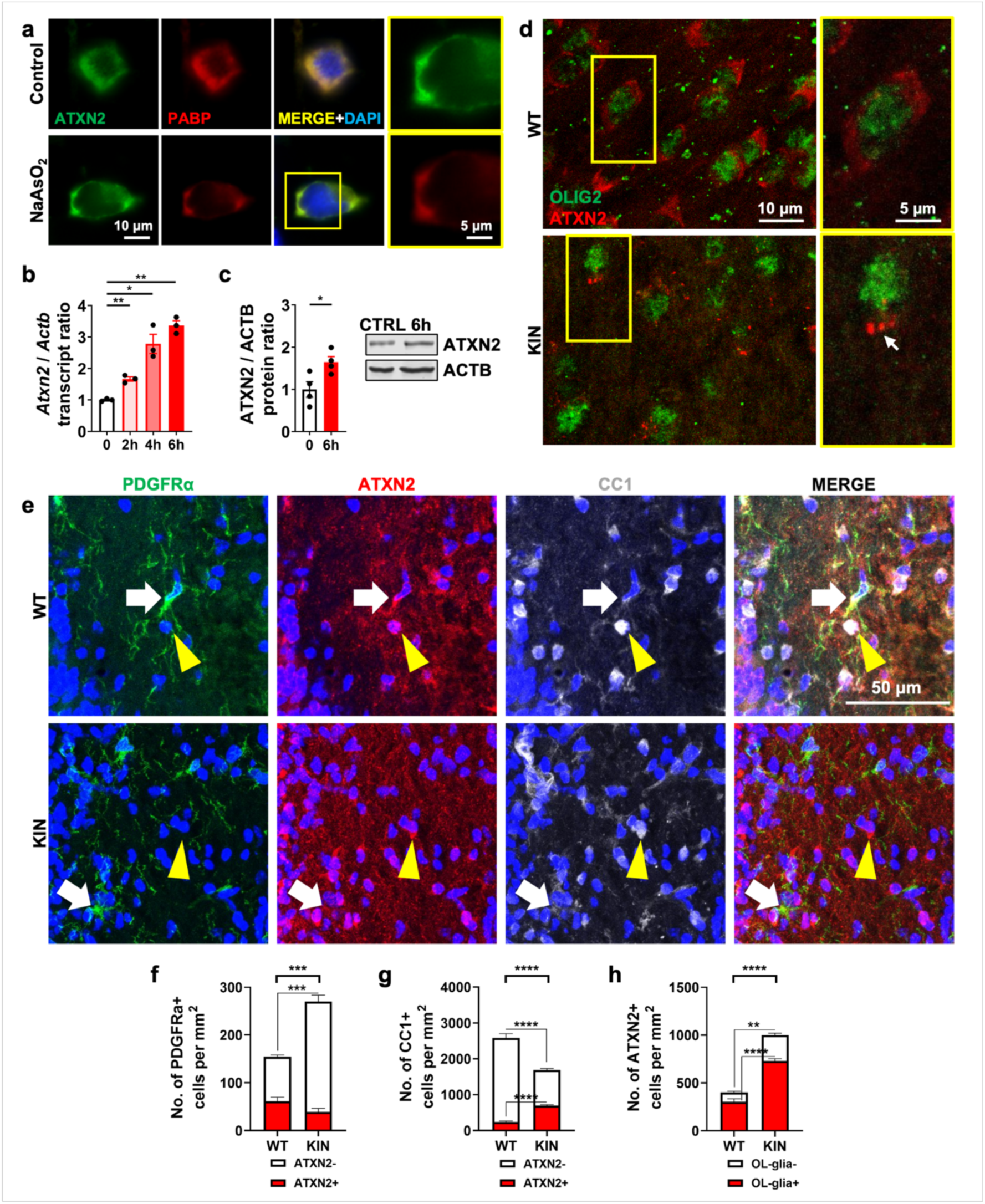
ATXN2 is expressed in oligodendrocytes and forms disease-associated aggregates. **a,** Expression and distribution of ATXN2 in oligodendrocytes was assessed using Oli-neu cells under normal and oxidative stress conditions. Normally diffuse ATXN2 (green) was found to relocate into SGs marked by PABP (red) under sodium-arsenite (NaAsO_2_) treatment. DAPI (blue) marks the nuclei. **b,** Expression of *Atxn2* transcript in differentiated Oli-neu cells was measured under normal growth conditions by qRT-PCR. Upon starvation stress, *Atxn2* transcript is significantly upregulated over time (2-6 h). *Actb* was used as housekeeping gene. Each data point represents a biological replicate. **c,** Presence of ATXN2 protein in differentiated Oli-neu cells was validated with quantitative immunoblots under normal growth conditions. ATXN2 protein abundance was significantly upregulated upon starvation stress at 6 h. ACTB was used as loading control. Each data point represents a biological replicate. **d,** Immunohistochemical assessment of ATXN2 protein (red) in OLIG2+ oligodendrocytes (green) in WT and KIN cerebella at terminal stage confirms its expression and diffuse distribution in WT oligodendrocytes, and shows its toxic cytosolic aggregation in KIN oligodendrocytes. **e,** Immunohistochemical assessment of ATXN2 co-localization within PDGFRα- and CC1-positive cells in cerebellar WM showed its expression in both OPCs and mature oligodendrocytes. **f,** Quantification of all OPCs in cerebellar WM showed a significant increase in KIN mice, however a reduction in ATXN2-positive portion was observed. **g,** Quantification of all mature oligodendrocytes in cerebellar WM showed a significant decrease in KIN mice, however the number of ATXN2-positive portion was significantly increased. **h,** Comparison of ATXN2 signals from oligodendroglia versus other cells in cerebellar WM showed majority of ATXN2 signal to come from oligodendroglial lineage in both WT and KIN mice.

Overall, these data for the first time confirm the substantial mRNA expression and protein abundance of ATXN2 in myelinating cells, as well as the polyQ expansion triggered cytosolic ATXN2 aggregate formation in oligodendrocytes, as a relevant contributor to SCA2 pathology.

### Myelin anomalies with prominent MAG depletion occur at pre-onset stage

In order to better understand the demyelination process at the molecular level, label-free global proteome analyses were performed in KIN cerebellum and spinal cord tissues at the terminal stage, identifying 3593 and 3769 proteins, respectively (**Supp. Figure S4a**). Among the 2999 identified proteins in common (**Figure 5a**), those with less than 20% dysregulation difference between both tissues were identified to determine “commonly dysregulated proteins” in spinocerebellar tissue. Common upregulations mostly consisted of RNA-binding proteins (RBPs), possibly due to their sequestration in the cytosolic ATXN2 aggregates, as a well-known phenomenon in this form of neuropathology. On the other hand, common downregulations mainly consisted of metabolic factors and myelin-associated proteins, representing previously understudied outcomes of ATXN2 pathology. Interaction network and pathway enrichment analyses of these proteins (with <70% abundance) revealed “Myelin sheath” as the most strongly downregulated entity after the general term “Cytoplasm” (**Supp. Table S1, Figure 5b**, marked with shades of purple). Besides myelin proteins, several synaptic and axonal proteins were also found downregulated in KIN spinocerebellar proteome (**Figure 5b**, marked with shades of orange and blue), including the two neurofilament heavy/light chain subunits NEFH/L, prompting us to characterize the temporal dynamics of neuronal vs. oligodendroglial marker deficits throughout the disease course.

**Fig. 5:**
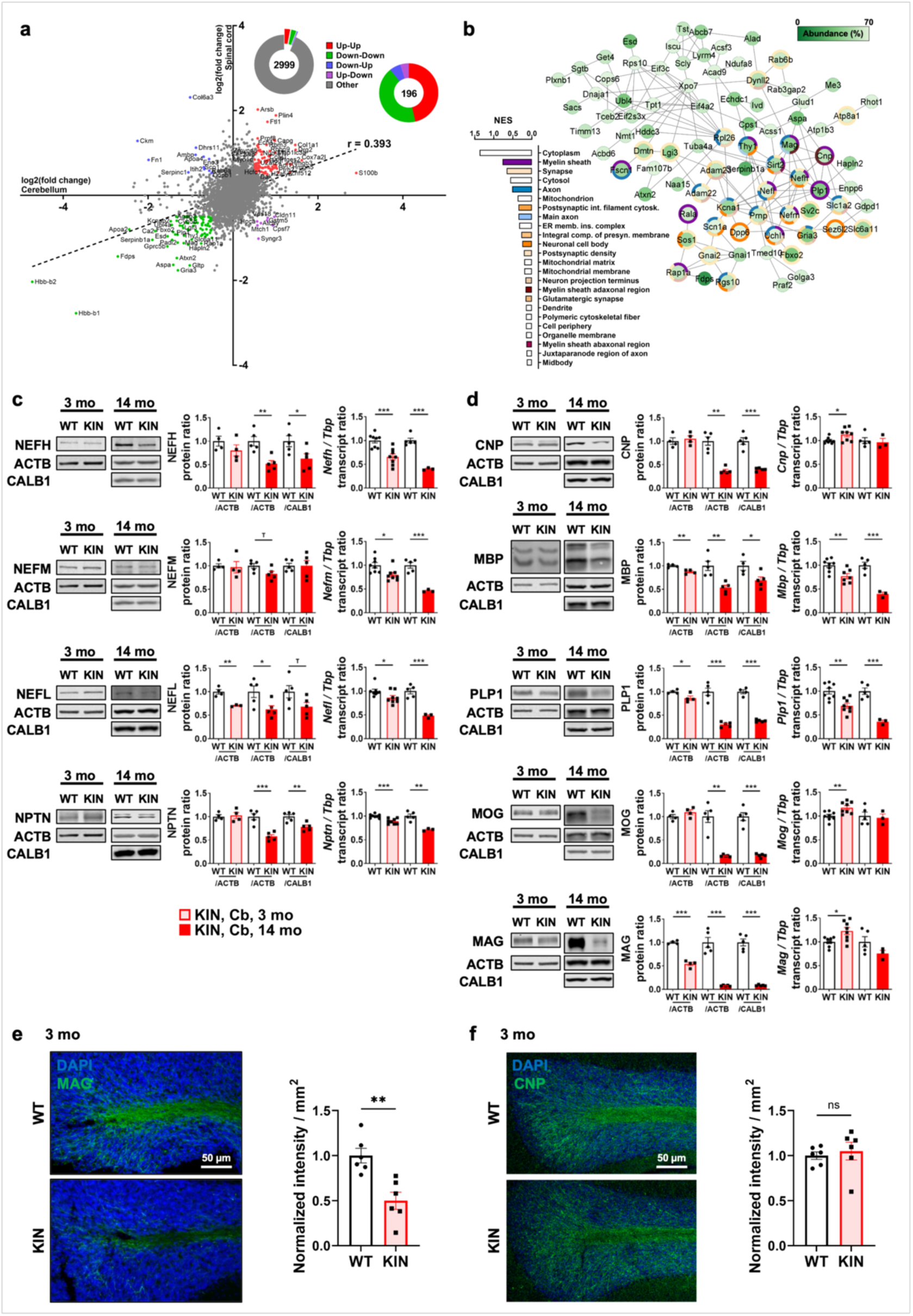
ATXN2 polyQ expansion toxicity progressively impairs myelin proteins in spino-cerebellar tissue. **a,** Graph depicting the dysregulation rates of commonly identified proteins in cerebellum and spinal cord samples from WT and KIN animals at terminal stage of 14 months old (r = 0.355, Pearson correlation). Pie charts depict the total number of commonly identified proteins (2999), all dysregulated proteins with >0.5 log_2_(fold change) difference in all directions (196), among which there were 91 proteins commonly upregulated (red), 84 proteins commonly downregulated (green), 12 proteins down in cerebellum, up in spinal cord (blue), and 9 proteins up in cerebellum, down in spinal cord (purple). **b,** Protein-protein interaction network and pathway enrichment analysis of the commonly downregulated proteins (with <70% abundance in KIN) showed a cluster of multiple myelin (in shades of purple) and axonal/synaptic proteins (in shades of blue and orange), with the term “Myelin sheath” appearing as the strongest dysregulation after the term “Cytoplasm”. **c,** Protein and transcript levels of neuronal compartment factors in WT and KIN cerebellum (Cb) at pre-onset (3 mo) and terminal (14 mo) stages, showing late onset downregulations for NEFH and NPTN, with an earlier downregulation of NEFL protein. ACTB was used as loading control, and normalizations against CALB1 were performed in order to test the neuropil affection versus Purkinje cell soma. All transcripts showed a significant downregulation at pre-onset stage with progressive reduction during disease course in KIN tissue. *Tbp* was used as housekeeping gene in qRT-PCR experiments. Each data point represents a single animal. **d,** Protein and transcript levels of myelin compartment factors in WT and KIN cerebellum at pre-onset (3 mo) and terminal (14 mo) stages, showing prominent downregulations for CNP, MBP and PLP1, with a massive reduction of MOG and MAG proteins. Slight yet significant reductions were observed in MBP and PLP1 levels at the pre-onset stage, accompanying a stronger downregulation of MAG. ACTB was used as loading control, and normalizations against CALB1 were performed in order to test the myelin affection versus Purkinje cell soma. *Mbp* and *Plp1* transcripts showed a progressive downregulation throughout the disease course in KIN tissue, however *Cnp*, *Mog* and *Mag* transcripts interestingly showed a significant upregulation at the pre-onset stage which disappeared at the terminal stage. *Tbp* was used as housekeeping gene in qRT-PCR experiments. Each data point represents a single animal. Immunohistochemical assessment and quantification of MAG **(e)** and CNP **(e)** proteins in WT and KIN cerebellum at pre-onset stage confirms the immunoblot findings with a prominent reduction of MAG abundance and no change in CNP levels in KIN tissue. At least two sections were quantified from two WT and KIN animals. DAPI (blue) marks the nuclei in granular layer.

To evaluate which of the neuronal vs. oligodendroglial anomalies are detectable already at the pre-onset stage, we assessed the transcript and protein levels of strong downregulations observed in proteome data. Both the somatodendritic marker CALB1 and the dendritic microtubule component TUBA4A showed minor reductions both at pre-onset and terminal stages, although their transcripts showed a progressive reduction over the disease course (**Supp. Figure S4b-e**), indicating a negligible loss of Purkinje neuron somata in KIN cerebellum even in terminal disease stage. Still, we decided to employ the neuronal marker CALB1 on the one hand, and the ubiquitous cell cytoskeleton component β-Actin (ACTB) on the other hand as loading control and normalizers at the terminal stage. Normalization against CALB1 would specifically reveal the relative dysregulation of query proteins with respect to Purkinje neuron affection. As CALB1 levels were found stable in pre-onset stage, this normalization was not performed.

Among the three neurofilament subunits, only NEFL showed a significant reduction both at pre-onset (3 months) and terminal (14 months) stages, while NEFH showed a significant reduction only at the terminal stage, and NEFM was found unaltered throughout the disease course (**Figure 5c**). Similar to NEFH, the synaptic adhesion marker neuroplastin (NPTN), which is important for the localization of GABA_A_ and GluA1 neurotransmitter receptors^48^, also showed a significant reduction only at terminal stage. Interestingly, all their transcript levels were found significantly downregulated already at pre-onset stage, decreasing further with disease progression.

Strong downregulations were confirmed at terminal stage also for the myelin membrane compaction regulators CNP and MBP, as well as the multilamellar myelin sheath maintenance factor PLP1, with both MBP and PLP1 showing mild downregulations already at the pre-onset stage (**Figure 5d**). Extreme reductions were observed at terminal stage for two myelin glycoproteins in axonal contact, MOG and MAG. MAG also exhibited a significant reduction at pre-onset stage, stronger than all the other proteins investigated thus far. This finding was confirmed by immunostaining WT and KIN cerebella at pre-onset stage: a strong reduction in MAG signal was evident in KIN tissue (**Figure 5e**), while CNP levels appeared comparable between WT and KIN samples (**Figure 5f**), as expected from the immunoblot findings. Although the transcript levels of *Mbp* and *Plp1* followed a similar pattern as neurofilament subunits with early and progressive downregulation throughout the disease course, no reduction was observed in the transcript levels of *Cnp*, *Mog* or *Mag* at the terminal stage. Unexpectedly, all their transcript levels were found significantly upregulated at the pre-onset stage (**Figure 5d**), possibly to compensate ATXN2-dependent defects.

A similar pattern of dysregulations was observed in the spinal cord at terminal stage, again with extreme downregulation of MAG protein with an effect size that exceeded the downregulation of NEFH as a prominent neuronal marker (**Supp. Figure S5a**). Immunohistochemical assessment of MAG protein in spinal cord sections confirmed these findings (**Supp. Figure S5b**). Further exploration of the neuronal and oligodendroglial proteins with strong downregulation in the proteome survey substantiated early significant and progressive mRNA reductions for the Internexin neuronal intermediate filament alpha (*Ina*), the Myelin-associated oligodendrocyte basic protein (*Mobp*), and a neuron-secreted factor of perineuronal networks and Ranvier nodes (*Hapln4*). Late-onset mRNA reductions were documented for the Myelin and lymphocyte protein (*Mal*) and the axon-axon adhesion repressor Reticulon-4 / Nogo (*Rtn4*), as well as three other components of the perineuronal networks and Ranvier nodes between myelinated axon segments (*Hapln1-3*), which control the diffusion of cations that are crucial for saltatory conductivity^49–51^ (**Supp. Figure S5c**).

Overall, the global proteome profile together with the targeted validations point to an exceptional disturbance of the cytoskeleton and adhesion factors, affecting oligodendroglia at least as early and strongly as neurons. Despite the prominent impacts of ATXN2 pathology being clarified in detail by these findings, the question still remained whether the axon-myelin disconnection was triggered by neuronal atrophy or oligodendroglial pathology.

### Alteration of myelin compartment factors does not occur as a secondary consequence to neuronal atrophy

To answer the question if ATXN2 polyQ expansion toxicity in neurons could trigger myelin damage as a secondary consequence, we examined a transgenic (Tg) SCA2 mouse model where human *ATXN2*-Q58 is overexpressed specifically in cerebellar Purkinje neurons under control of the *Pcp2* promoter^38^. Previous reports showed these mice to lose more than half of their cerebellar Purkinje neurons at the late-symptomatic stage of 46 weeks evidenced by CALB1 immunohistochemistry^38^. However here, we found only a minor downregulation of *Mag*, *Mog* and *Mbp* in *ATXN2*-Q58-Tg cerebella solely at the late-symptomatic stage, with no dysregulations at pre-onset stage of 16-weeks (**Supp. Figure S6**). We previously reported a profound pre-onset deficit of NAT8L in the KIN cerebellum, a neuronal enzyme that provides the metabolic building block N-acetylaspartate (NAA) to oligodendrocytes for myelination. The very subtle affection of *Nat8l* in *ATXN2*-Q58-Tg cerebella only at the late-symptomatic stage, and the complete lack of its dysregulation at pre-onset stage showed good agreement with other myelin factors in this model. Therefore, an expansion size of Q58 in transgenic mice overexpressing ATXN2 only in cerebellar Purkinje cells seems to preferentially affect neuronal perikarya. In comparison, larger expansion size of Q100 in our KIN mouse that expresses ATXN2 at endogenous levels in all cell types appears to affect the neuropil much more, particularly the myelinated axons. Thus, the motor deficits of KIN mice are largely due to dysregulated neural connections rather than neuronal cell death, a phenomenon that was not observed even at the terminal disease stage. Collectively, these data indicate that the massive neurotoxicity of transgenic ATXN2 expansion specifically in neurons could only trigger very mild transcriptional dysregulations in oligodendrocytes, arguing against the notion of demyelination in KIN mice to be secondary to neuronal pathology.

### ATXN2 polyQ expansion toxicity impairs alternative splicing of *Mag* and *Plp1*

In neurons, ATXN2 interacts with various known splice factors, *e.g.* TDP-43 and A2BP1/RBFOX1, which control neuronal excitation. While no investigation so far has characterized the interactors of ATXN2 in oligodendroglia, we noticed that several of the dysregulated myelin proteins have multiple splice isoforms. MAG is encoded by large and small isoforms of *Mag* transcript (*L/S-Mag*, with the large isoform expressed during development), and PLP1 is encoded by long *Plp1* or shorter *Dm-20* which lacks exon IIIb (*Plp1* being expressed in oligodendrocyte progenitors)^52–54^. Both these splicing events are performed by the same RBP named Quaking (QKI), the loss of which leads to relative increase of *S-Mag* and *Dm-20* isoforms as depicted in **Figure 6a, c**^55,56^. Assessment of *Mag* and *Plp1* splicing in KIN mouse tissues with semi-qPCR revealed a dramatic dysregulation especially in the terminal stage, with a milder yet significant impact in the pre-onset stage for both transcripts. Progressive reduction of *L-Mag* in cerebellum and spinal cord contrasted with an increase in *S-Mag* levels (**Figure 6b**). In addition, strong deficits of both *Plp1* isoforms (*Plp1* and *Dm-20*) were observed, again with preponderance of the mature isoform in old tissues (**Figure 6d**). Thus, the commonly used measures of *Mag* and *Plp1* splicing, i.e. *L-/S- Mag* and *Dm-20/Plp1* ratios, demonstrated the progressive nature of splicing defects. These findings were reproduced independently with different set of primers designed for qRT-PCR (**Supp. Figure S7a**). The lack of splicing dysregulations in *Atxn2* knock-out (KO) cerebellum demonstrated their specificity for the ATXN2 polyQ expansion pathology (**Figure 6b, d, Supp. Figure S7b**). Analysis of the *ATXN2*-Q58-Tg cerebellum at two ages did not reveal any major splice defects (**Supp. Figure S7c**), once again indicating that the oligodendroglia abnormalities are not secondary to neurotoxicity. In contrast to *Mag* and *Plp1*, *Cnp* pre-mRNA is alternatively spliced by QKI-independent mechanisms, although its primary splice factors remain unknown^57^. Examination of *Cnp* variants revealed no dysregulation in KIN cerebellum at terminal stage (**Supp. Figure S8**), and overall unchanged levels of all isoforms agreed with the previous assessment of its total levels presented in Figure 5d. Strong and progressive dysregulation of *Mag* and *Plp1* splicing, next to lack of any splice irregularity in *Cnp* variants, suggest a selective involvement of ATXN2 toxicity in their splicing via QKI.

**Fig. 6:**
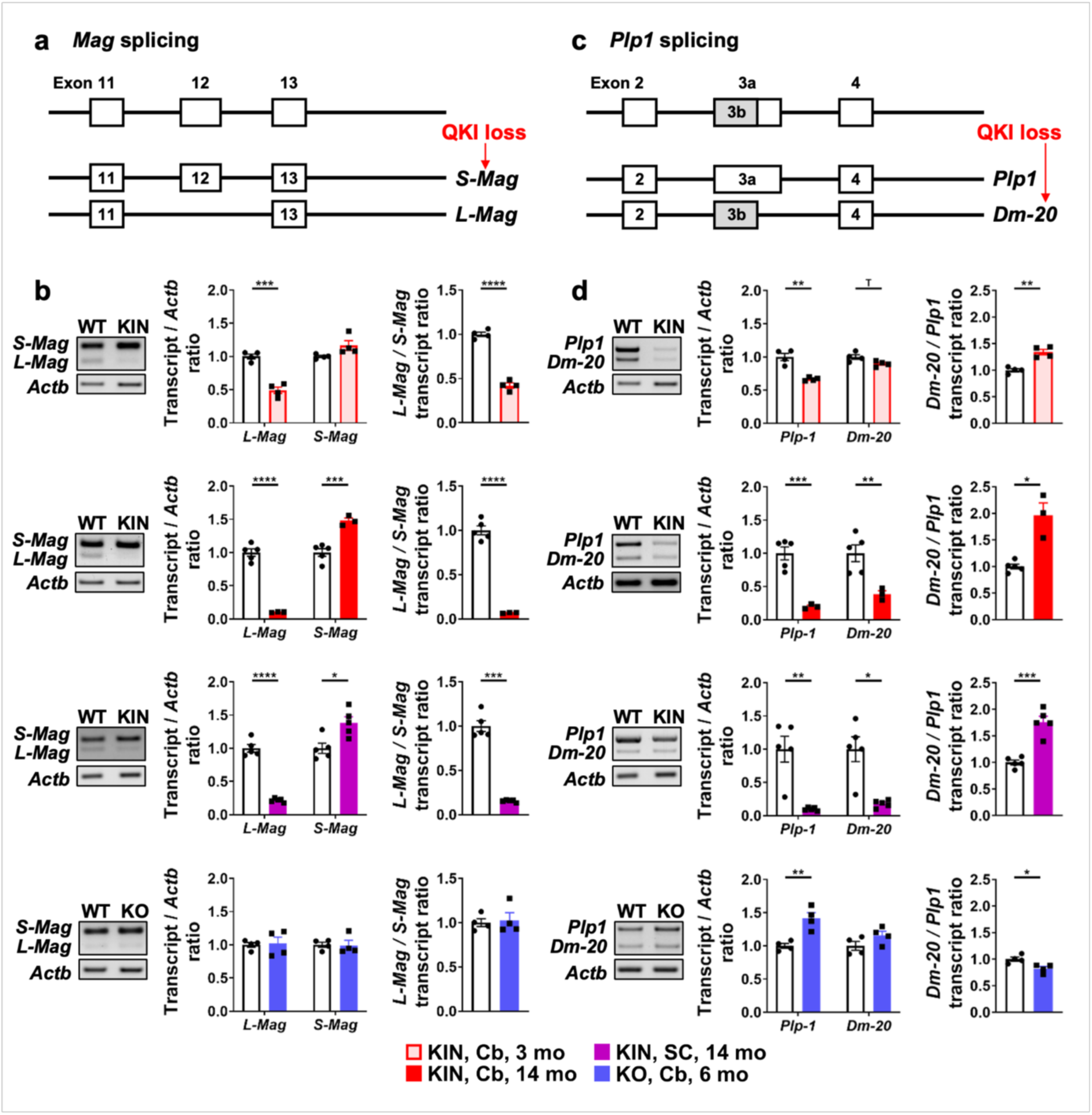
ATXN2 aggregation disrupts the alternative splicing of mature myelin markers *Mag* and *Plp1*. **a, c,** Schemes depicting the alternative splicing patterns of *Mag* and *Plp1* transcripts, with indication of which isoform gets upregulated in the case of QKI loss. Expression levels of *Mag* **(b)** and *Plp1* **(d)** splice isoforms were measured by semi-qPCR in KIN cerebellum (Cb, 3 mo, 14 mo), KIN spinal cord (SC, 14 mo) and KO cerebellum (Cb, 6 mo) in comparison to age-matched WT controls. Strong splicing disturbances indicated by *L-Mag/S-Mag* and *Dm-20/Plp1* ratios were observed readily at pre-onset stage (3 mo) in KIN cerebellum, that showed further progression until the terminal stage (14 mo) in both cerebellum and spinal cord. Lack of a similar missplicing pattern in KO cerebellum ensured the ATXN2 aggregation-dependent nature of these dysregulations. The mild alteration of *Plp1* splicing in KO samples observed here was not confirmed by further qRT-PCR experiments (see **Figure S7b**). *Actb* was used as housekeeping gene and loading control in semi-qPCR experiments. Each data point represents a single animal.

### Loss of proper QKI function explains the alternative splicing defects of *Mag* and *Plp1*

The genetic deletion of QKI in mouse causes massive reduction of the developmental alleles, *L- Mag* and *Plp1*^55,56^, as observed in KIN tissue. QKI belongs to the KH-family RBPs, other members of which include the nuclear SAM68 protein, previously observed to respond to ATXN2 polyQ expansion toxicity in cerebellar neurons with elevated abundance^58^. Distinct QKI isoforms predominantly localize to the nucleus (QKI5), to the cytoplasm (QKI7), or both (QKI6)^59–61^.

The subcellular distribution of QKI proteins, as well as their association with ATXN2 in SGs were validated in cultured Oli-neu cells. Under normal growth conditions, QKI5 displayed a dominantly nuclear localization, whereas QKI6 and QKI7 were observed in both cytosolic and nuclear compartments (**Figure 7a-c**). Under oxidative stress, QKI6 showed the highest rate of translocation into cytosol and the strongest association with ATXN2 in SGs, although a portion of cytosolic QKI7 and a lesser portion of QKI5 also displayed association with ATXN2 in SGs.

**Fig. 7:**
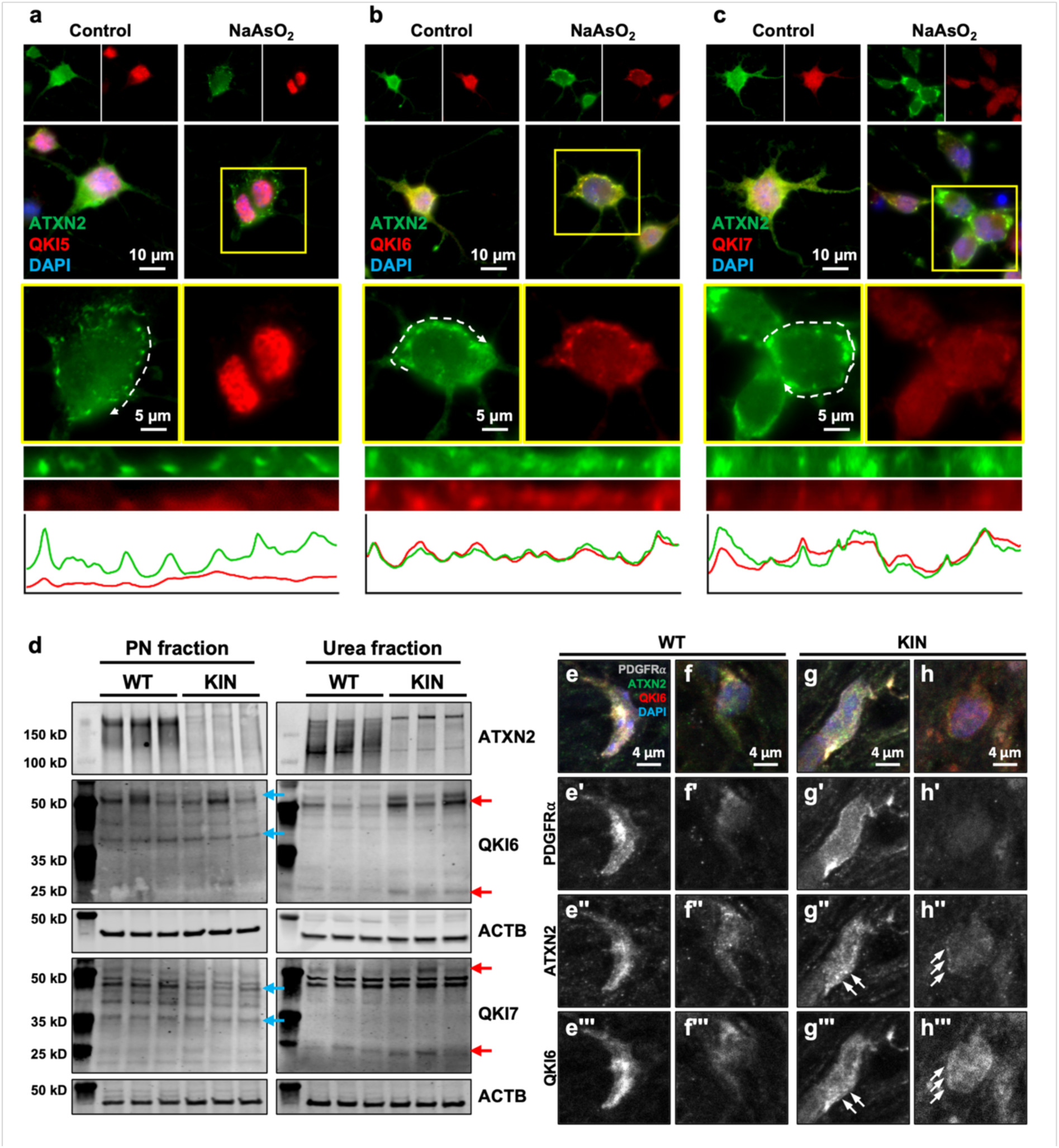
Mutant ATXN2 sequesters QKI into pathological aggregates. **a-c,** Subcellular distribution of QKI isoforms (red) and their co-localization with ATXN2 (green) were investigated using Oli-neu cells under normal and oxidative stress conditions. DAPI (blue) marks the nuclei. Under normal growth conditions, QKI5 showed a nuclear localization, while QKI6 and QKI7 were detected in both cytosol and nucleus. Under oxidative stress induced by sodium-arsenite (NaAsO_2_), QKI6 showed the highest rate of association with the SGs marked by ATXN2, although a portion of QKI7 and a smaller portion of QKI5 also co-localized with SGs. Insets marked with yellow squares in NaAsO_2_ images were shown enlarged in lower panels. White dotted lines mark the straightened segments shown below. Normalized plot profiles of each channel in straightened segments depict the co-localization of respective QKI isoform with ATXN2 in SGs. **d,** Separation of WT and KIN cerebella at the terminal stage into PN (soluble) and Urea (insoluble) fractions showed an increased abundance of QKI6 and QKI7 to co-fractionate with mutant ATXN2 in the insoluble fraction (red arrows). The segregation of both QKI isoforms at different levels than that of their original size (≈35 kDa) in Urea fraction is suspected to be the result of post-translational modifications they encounter at the aggregates. **e-h,** Immunohistochemical assessment of ATXN2 and QKI6 co-aggregation in PDGFRα- positive and negative cells of the white matter (OPCs and more mature stages of the lineage, respectively). At least two sections were imaged from three WT and KIN animals with similar findings.

When we analyzed the expression levels of these QKI family members in the terminal stage of KIN mice by quantitative immunoblots, we observed a decrease of the cytoplasmic isoforms QKI6 and QKI7 versus an increase of nuclear QKI5 (**Supp. Figure S9a**), well correlated with a previous report stating that this isoform bias in transcriptome is produced by QKI malfunction^59^. Among the three isoforms, cytoplasmic QKI6 changes from higher abundance at 3 months to a depletion at 14 months, and was previously observed to relocalize into SGs^61^, thus representing a perfect candidate to be sequestrated into ATXN2 aggregates. Indeed, immunohistochemical studies in terminal stage cerebellum demonstrated its sequestration into cytoplasmic aggregates marked by the ATXN2 interactor PABP (**Supp. Figure S9b**). In order to separate soluble ATXN2 from insoluble aggregates formed by Q100-expanded protein, and to analyze QKI isoforms in these fractions, cerebella at terminal stage were sequentially lysed in NP-40/PBS (PN) and Urea-based buffers. The majority of wild-type ATXN2 appeared in “soluble” PN fractions, while the majority of normally low-abundance ATXN2-Q100 appeared in “insoluble” Urea fractions, as expected (**Figure 7d**). Multiple bands were observed for both QKI6 and QKI7 in soluble fractions in WT and KIN samples, both at their expected size of ∼35 kD and higher around 50 kD (**Figure 7d**, blue arrows). A reduction in the abundance of all QKI7 bands were obvious, however there was no striking difference in QKI6 abundance in the soluble fraction. Notably, QKI6 and QKI7 immunoreactivity in insoluble fractions appeared at 25 kD and 50 kD (**Figure 7d**, red arrows), and showed visibly higher levels in KIN samples suggesting a strong interaction with mutant ATXN2. RBPs are often modified in pathological aggregates mainly by phosphorylation, ubiquitination or proteolytic cleavage, which causes them to migrate at unexpected sizes in an acrylamide gel^62,63^. We believe these, yet unidentified, modifications underlie our observation of QKI6 and QKI7 immunoreactivity at 25 and 50 kD levels.

To further dissect whether ATXN2 and QKI6/QKI7 normally interact or aggregate in OL cells, we assessed their localization together with PDGFRα staining in WT and KIN cerebella. While we did not manage to get a reliable result from QKI7 immunohistochemistry, in PDGFRα-positive OPCs, both ATXN2 and QKI6 reliably displayed cytosolic distributions in WT mice (**Figure 7e**) and accumulated into small puncta in KIN samples (**Figure 7g**). In PDGFRα-negative, but QKI6- positive cells of the white matter indicating later stages of oligodendroglial lineage, both ATXN2 and QKI6 abundance was much lower, yet still showed a relatively cytosolic distribution in WT (**Figure 7f**) and multiple small aggregates in KIN samples where QKI6 seemed to colocalize with mutant ATXN2 (**Figure 7h**).

Examining the transcript levels of *Qki* splice isoforms documented tissue-specific dysregulations only at the terminal stage, suggesting their secondary nature to pre-onset protein alterations. Increased levels of *Qki5* and *Qki6* with concomitant *Qki7* deficit were observed in cerebellum. In the more strongly affected spinal cord, both *Qki6* and *Qki7* transcripts were decreased (**Supp. Figure S9c**). Again, such isoform bias was observed neither in *Atxn2*-KO cerebellum (**Supp. Figure S9d**), nor under selective neurotoxicity in *ATXN2*-Q58-Tg cerebellum (**Supp. Figure S9e**). Overall, the data indicate that the missplicing of myelin proteins occurs due to functional depletion of QKI via its sequestration into protein aggregates both in earlier and later stages of oligodendroglial lineage.

### Purkinje cell responses to whisker stimulation are delayed by ATXN2 polyQ expansion

One of the main functional roles of myelin layers surrounding axons is to increase the signal speed in neuronal circuits. Balanced conduction speed is of great relevance when monitoring the environment, responding to sensory stimuli and generating coordinated motor responses. To test the consequences of demyelination in KIN mice, we employed an established paradigm that documents Purkinje cell responses to whisker stimulation^64–66^. A stimulus to the mouse whiskers travels via the trigeminal nerve directly to the inferior olive, where climbing fibers originate and have therefore high chance to generate complex spikes in Purkinje cells located in lobules crus 1 and crus 2 with a latency of approximately 45 ms (**Figure 8a, b**). Given the simplicity of this tri- synaptic circuit, any delay in the complex spike modulation to the whisker stimulation would unmask a deficit in the conduction speed of those neurons. Under our experimental conditions, Purkinje cells recorded from KIN mice showed responses in complex spike at a significantly larger latency (KIN 59±16 ms vs WT 45±19 ms, p=0.001) but similar in amplitude to the ones recorded from WT mice (p=0.226) after air-puff stimulation (**Figure 8c-g**). Overall, our data demonstrate significantly reduced speed conductance within the trigeminal circuit of KIN mice that can be explained by the demyelination.

**Fig. 8:**
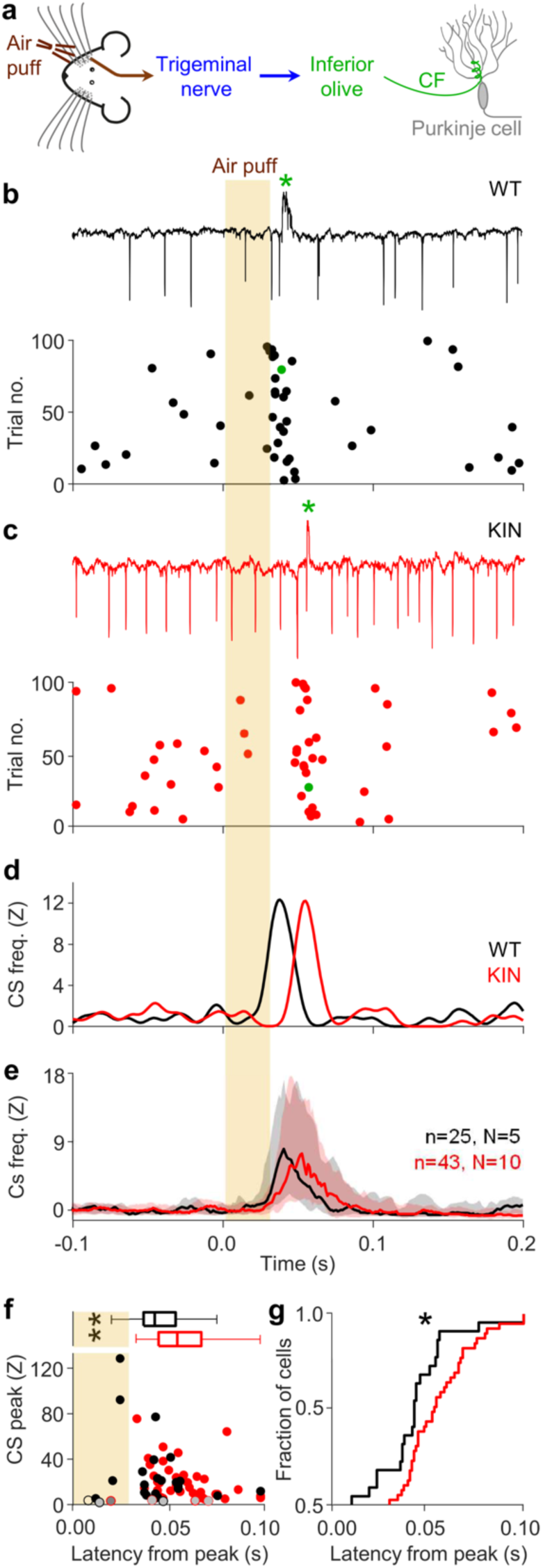
Delayed sensory responses in Purkinje cells of *Atxn2*-CAG100-KIN mice. **a,** Air puff stimulation of the whisker pad activates neurons in the sensory trigeminal nuclei (TN). From the TN, there is a direct projection to the inferior olive (IO), from where climbing fibers (CF) convey the sensory signal to Purkinje cells in the cerebellum. **b**, Extracellular recording of a representative Purkinje cell in an awake WT mouse. Top: single trace, with the green asterisk indicating the occurrence of a complex spike. Bottom: raster plot showing the distribution of complex spikes during 100 consecutive trials. The shaded area indicates the timing of the air puff. **c**, Same as b, but from a representative Purkinje cell in a KIN mouse. **d**, Convoluted peri-stimulus time histograms showing the complex spikes responses of the two example cells in b and c (see Methods). **e**, Median values of 25 WT and 43 KIN Purkinje cells. Shaded areas show the interquartile ranges. **f**, For each Purkinje cell, the amplitudes and latency of the maximum complex spike firing within the first 100 ms after whisker pad stimulation. Purkinje cells without a statistically significant complex spike response (Z < 3) are indicated with a grey filling, and they are excluded from further analysis. Purkinje cells recorded from KIN mice show significantly delayed complex spike responses (t test, p = 0.007, t = 2.847, df = 39). **g**, Cumulative distributions of the latencies of maximum modulation for WT (black) and KIN (red) Purkinje cells (two-sample Kolmogorov-Smirnov test, p = 0.011).

## Discussion

Current concepts of SCAs and ALS are focused on the neurodegenerative process, viewing glia responses as secondary consequence of compensatory character in the case of astrogliosis and microgliosis, but the affection of oligodendroglia and myelin is poorly understood. This study characterized the cerebellar and cerebral demyelination that accompanies the neuropil degeneration in ATXN2 expansion pathology, defining an underlying oligodendroglia maturation problem with prominent depletion of myelin proteins. We found that ATXN2 polyQ expansion leads to QKI sequestration into cytosolic aggregates within oligodendrocytes, therefore leading to its partial loss of function and causing missplicing of *Mag* and *Plp1* transcripts, demonstrating for the first time an oligodendroglial component in SCA2 pathogenesis. Our findings document the white matter reduction in KIN cerebellum exceeding gray matter reduction (**Figure 1**), with clear demyelination on TEM imaging and axonal sprouting as a result, which was also observed in a SCA2 patient sample (**Figure 2**). Detailed quantification of oligodendroglia lineage revealed an increased number of proliferating OPCs both immediately after, and following a wash-off period after BrdU administration in KIN mice, suggesting an urge to restore the myelin loss. While the mature oligodendrocytes also showed an increased number in KIN mice immediately after BrdU treatment, they failed to persist until the end of the wash-off period, indicating the interference of ATXN2 toxicity at this stage (**Figure 3**). We show for the first time that ATXN2 is also expressed in oligodendrocytes, and forms disease-associated aggregates. In fact, the majority of ATXN2 signals in cerebellar WM was found in oligodendrocyte lineage cells among all the glia in both WT and KIN animals (**Figure 4**). Further biochemical analyses revealed prominent downregulation of myelin compartment proteins in spinocerebellar proteome of KIN mice (**Figure 5**), along with significant alterations in alternative splicing of key myelin proteins (**Figure 6**), which can be attributed to the sequestration of QKI as their splicing factor into ATXN2-driven aggregates (**Figure 7**). The impact of these molecular findings was confirmed at the organism level by electrophysiological recordings of Purkinje cell response to sensory stimulation. We observed that the sensory-evoked Purkinje cell complex spike responses were equally strong, but delayed in KIN mice, suggesting a decreased axonal conduction speed (**Figure 8**) as a likely result of demyelination^67–69^, and as also observed in patients with SCA2^70,71^.

In conjunction with past reports, the following scenario of pathogenesis seems plausible. Previous work showed SAM68 to be a key factor in the degeneration of cerebellar neuron projections in SCA2^58^, and the present work identifies QKI as a key factor in the oligodendroglia pathology. Both proteins are part of the STAR family of RBPs with one KH domain, so it is important to consider their cellular roles and depict the cascade of molecular events to better understand the ATXN2 polyQ expansion pathology in both neurons and oligodendroglia. A part of the ATXN2 pool associates with SH3 domain proteins such as GRB2, SRC or FYN in the receptor tyrosine kinase endocytosis machinery, modulating the uptake of trophic signals and nutrients^17,72,73^. In neurons, GRB2/SRC control SAM68 relocalization to the nucleus^74^, while in oligodendrocytes FYN controls QKI function through its phosphorylation^75^. The association with SAM68 and subsequent phosphorylation modulate its binding to RNA and splicing activity in neurons^76^. In oligodendrocytes, QKI function is crucial for the formation and maintenance of axoglial junctions, and its depletion impairs axon myelination that leads to ataxia in mice^77^. Our findings of impaired QKI-mediated splicing, and consequent loss of MAG and PLP1 proteins reflect the central role of QKI in myelin maintenance as described before.

The ATXN2 ortholog in *C. elegans* (Atx-2) promotes germline differentiation through trophic signaling pathways that regulate mRNA translation and processing. This pathway involves two KH-domain RBPs, including the QKI ortholog GLD-1^78^. GLD-1 cooperates with CGH-1 (ortholog of the ATXN2-interactor DDX6) in regulating mRNA storage, and with ASD-1 (ortholog of the ATXN2-interactor A2BP1/RBFOX1) to control RNAs that regulate development^79^. In agreement, our data support QKI being a crucial transducer of trophic signals to oligodendrocyte maturation, a process disrupted in SCA2 due to its sequestration into ATXN2 aggregates. This evolutionary conservation underlines the fundamental nature of ATXN2-QKI interactions. The partial loss-of-function of QKI and other STAR family members, such as SAM68 that modulates mTOR pathway – a key regulator of myelination^80^, and synaptic plasticity in glutamatergic and GABAergic neurons^81^ – likely contributes not only to the loss of myelin lipids and cerebellar neuropil, but also to the loss of brain and body weight previously described in SCA2.

Among the molecular events downstream of QKI sequestration, the massive loss of the *L-Mag* splice variant and of MAG protein (<20%) was prominent. MAG and Nogo-A (encoded by *Rtn4*) interact as alternative ligands for the receptor NgR1, which modulates neurite outgrowth. Interestingly, Nogo-A was also transcriptionally downregulated by ATXN2 polyQ expansion toxicity. In addition, a previous report showed Nogo-A knock-out to trigger a transcriptional upregulation of *Atxn2* and *Atxn1* in mouse^82^, so this axon-myelin adhesion disturbance might be among the important primary events of ataxia pathogenesis. Actually, several reports recently showed the significance of early oligodendrocyte dysfunction preceding onset of disease signs in other polyQ-related neurodegenerative disorders, such as SCA1, SCA3, and Huntington’s disease^83–85^, which aligns with our findings here for SCA2.

In view of reported motor neuron disease and SCA1 patients with anti-MAG autoantibodies^86–88^, it is possible that aberrant splicing leads to MAG misfolding and autoantibody generation as a consequence of ATXN2 expansion. Genetic evidence also points to MAG as a crucial factor in the pathogenesis of ataxias, since three independent reports so far described families with MAG mutations to display corticospinal and cerebellar degeneration^89–91^, further connecting our findings to broader ataxia syndromes. In comparison, mutations in PLP1 also cause ataxia, but considerably more spasticity^92^. In view of the important role of MAG as a common target of autoimmune attack in multiple sclerosis^93^, it is conceivable that genetic factors like ATXN2 might play a exacerbating or protective role in some variants of multiple sclerosis. Indeed, such evidence was recently reported although the exacerbation effect of ATXN2 expansion was not general or strong^94,95^.

While this study provides the first description of oligodendroglial pathology in SCA2, future work should address the cell-autonomy of ATXN2 toxicity in oligodendrocytes by selective expression of the polyQ-expanded protein in these cells, or by its selective elimination in the otherwise KIN background. Both approaches have been used previously in Huntington’s disease to reveal the cell-autonomous toxicity of polyQ-expanded Huntingtin protein^85,96,97^. Single-cell transcriptomic and proteomic approaches would also shed further light onto ATXN2 toxicity-driven dysregulations in each cell type, not only providing new pathological hallmarks of the disease, but also pointing towards novel therapeutic opportunities. Indeed, the use of such technologies in SCA1 and SCA7 has recently identified new potential disease drivers and preferential affection of certain Purkinje cell subtypes, respectively^84,98^.

## Conclusions

This study used gross brain morphology, cell markers, and ultrastructural evidence to document impaired oligodendroglial maintenance and demyelination due to ATXN2 polyQ-expansion pathology, occurring via failed maturation of myelin upon missplicing of key structural proteins. Significant reduction in trigeminal circuit conduction speed demonstrated the functional consequence of this demyelination. The oligodendroglial splicing factor QKI is central to these defects, being sequestered into ATXN2-driven aggregates and undergoing a partial loss-of-function within oligodendrocytes (**Figure 9**). The main downstream effects are the depletion of the myelin glycoproteins MAG and MOG, which are also main attack sites for autoimmune demyelinating processes. These glial data and previous observations on neuronal SAM68 dysregulation reveal a consistently prominent pathogenic role of the STAR protein family for RNA-processing pathology in neurodegenerative diseases. Overall, this pioneer description of extra-neuronal ATXN2 toxicity supports the emerging gliocentric perspective in neurodegenerative research, as previously shown for other neuron-related diseases like autism-spectrum disorders, SCA1, SCA3 and Huntington’s disease^83–85,96,99^.These insights also raise hope that not only ALS but also other neuron/myelin disorders triggered within this pathway may be amenable to preventive therapy by ATXN2 depletion.

**Fig. 9:**
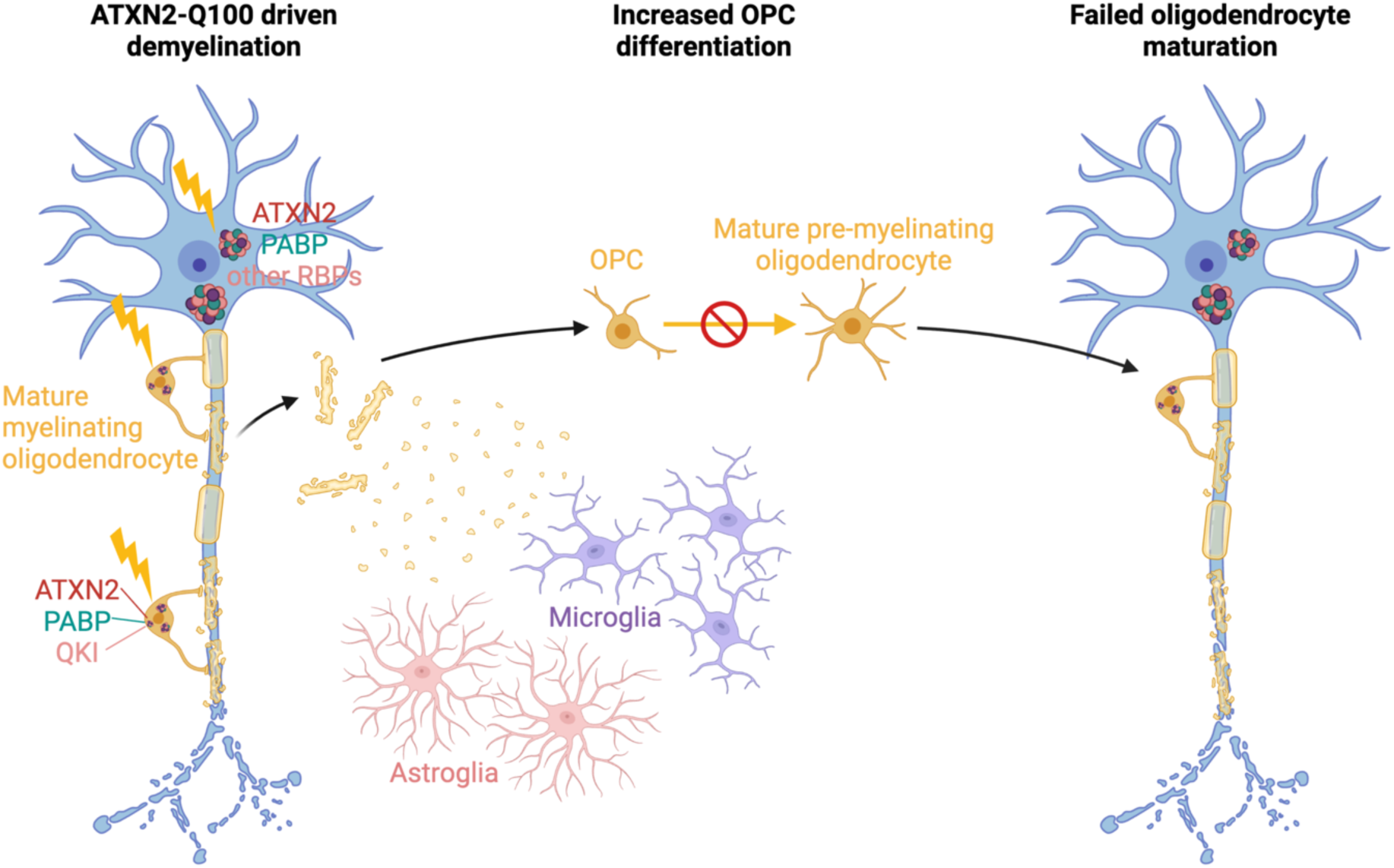
Schematic summary of ATXN2 pathology in neurons and oligodendrocytes. Progressive aggregation of ATXN2 in neurons is a hallmark of its toxicity across multiple neurodegeneration syndromes. These aggregates also sequester other RNA-binding proteins, including PABP, TDP-43, and FUS, disrupting a multitude of subcellular processes ranging from altered autophagy to axonal cargo transport, which ultimately lead to the degeneration of axon terminals and cell somata. Our findings from the *Atxn2*-CAG100-KIN mouse that expresses the mutant protein throughout the body at an endogenous level show that Purkinje cell somata are largely preserved even at the terminal stages of the disease course, whereas demyelination, and consequently, axon-myelin disconnection are among the first events of pathogenesis. Damaged myelin stimulates OPC proliferation and differentiation, however with unsuccessful maturation. We show that ATXN2 aggregates also form in cerebellar oligodendrocytes, sequestering the splice factor QKI, which leads to a dramatic loss of key mature myelin proteins, such as L-Mag and Plp1. Additionally, pronounced astrogliosis and microgliosis are also observed in KIN tissue, likely as a consequence of myelin and axonal damage.

## List of Abbreviations

A2BP1: Ataxin-2-binding protein 1, RBFOX1
ACTB: Actin beta
ALS: Amyotrophic lateral sclerosis
ASD-1: Alternative splicing defective 1, A2BP1/RBFOX1 ortholog in C. elegans
ASPA: Aspartoacylase
Atx2: Ataxin-2 ortholog in C. elegans
Atxn1: Ataxin-1
ATXN2: Ataxin-2
BMI: Body mass index
BrdU: 5-bromo-2-deoxyuridine
CAG: Cytosine-Adenosine-Guanine
CALB1: Calbindin 1
CC1: Anti-adenomatous polyposis coli (APC) clone CC1
CGH-1: Conserved germline helicase 1, DDX6 ortholog in C. elegans
CNP: 2’,3’-Cyclic nucleotide 3’ phosphodiesterase
DAPI: 4′,6-diamidino-2-phenylindole
dbcAMP: Dibutyryl cyclic adenosine monophosphate
DDX6: DEAD-box helicase 6
Dm-20: Shorter splice variant of Plp1 pre-mRNA that lacks exon IIIb
ERGIC: Endoplasmic reticulum-Golgi intermediate compartment
FDR: False discovery rate
FUS: Fused in sarcoma
FYN: FYN proto-oncogene, Src family tyrosine kinase
GABA: Gamma-aminobutyric acid
GFAP: Glial fibrillary acidic protein
GLD-1: Defective in germ line development 1, QKI ortholog in C. elegans
GluA1: Glutamate ionotropic receptor AMPA type subunit 1
GM: Gray matter
GRB2: Growth factor receptor bound protein 2
Hapln1-4: Hyaluronan and proteoglycan link protein 1-4
IBA1: Allograft inflammatory factor 1, AIF1
Ina: Internexin neuronal intermediate filament alpha
KH: K-homology domain
Ki67: Marker of proliferation Ki-67
KIN: Knock-in
KO: Knock-out
LC: Liquid chromatography
LSm: Like-SM RNA binding domain
LSmAD: LSm-associated domain
MAG: Myelin associated glycoprotein
Mal: Myelin and lymphocyte protein
MAP-Tau: Microtubule associated protein Tau
MBP: Myelin Basic Protein
Mobp: Myelin-associated oligodendrocyte basic protein
MOG: Myelin oligodendrocyte glycoprotein
MS: Mass spectroscopy
MSA: Multiple system atrophy
mTOR: Mechanistic target of rapamycin
NAA: N-acetylaspartate
NaAsO2: Sodium arsenite
NAT8L: N-acetyltransferase 8 like
NEFH-M-L: Neurofilament heavy-medium-light chain
NG2: Chondroitin sulfate proteoglycan 4, CSPG4
NgR1: Nogo (Reticulon 4) receptor 1
Nogo-A: Reticulon 4, RTN4
NOTCH1: Notch receptor 1
NPTN: Neuroplastin
OL: Oligodendroglial lineage
OLIG2: Oligodendrocyte transcription factor 2
OPC: Oligodendrocyte precursor cell
Pab1: Poly(A) binding protein, PABP ortholog in S. cerevisiae
PABP: Poly(A)-binding protein
PAM2: PABP-interacting motif 2
Pbp1: Pab1-binding protein, ATXN2 ortholog in S. cerevisiae
Pcp2: Purkinje cell protein 2
PDGFR⍺: Platelet derived growth factor receptor alpha
PLP1: Proteolipid protein 1
polyQ: Poly-glutamine
Q: Glutamine
QKI: Quaking
RBFOX1: RNA binding Fox-1 homolog 1, A2BP1
RBP: RNA-binding protein
RT: Room temperature
RT-PCR: Reverse transcription PCR
Rtn4: Reticulon-4 / Nogo
SAM68: KH RNA binding domain containing, signal transduction associated 1
SCA2: Spinocerebellar ataxia type 2
SG: Stress granule
SH3: SRC homology 3 domain
SNCA: Synuclein alpha
SRC: Protooncogene SRC, Rous sarcoma
STAR: Signal transduction and activation of RNA family of proteins
TBP: TATA-Box binding protein
TDP-43: TAR DNA binding protein
TEM: Transmission electron microscopy
Tg: Transgenic
TUBA4A: Tubulin alpha 4a
UTR: Untranslated region
WM: White matter
WT: Wild type

## Declarations

### Ethics approval and consent to participate

Not applicable.

### Consent for publication

Not applicable.

### Availability of data and materials

The mass spectrometry data have been deposited to the ProteomeXchange Consortium (http://proteomecentral.proteomexchange.org) via the PRIDE partner repository^43^ with the dataset identifier PXD036124 (Project DOI: 10.6019/PXD036124).

### Authors’ contributions

All authors read and approved the final manuscript. Conceptualization: NES, JEVB, LD, GA, Data Curation: AJM, LB, VR, DM, Formal Analysis: NES, JEVB, JOA, AA, AJM, LB, VR, MP, DM, JK, MF, Funding Acquisition: GA, LD, DM, AG, TD, LWJB, CIDZ, Investigation: NES, JEVB, JOA, AA, JCP, KS, EF, ZEK, LEAM, Methodology: NES, JEVB, JOA, SG, LD, GA, Project Administration: LD, GA, Resources: JCP, MH, LAB, SG, AG, TD, LWJB, CIDZ, Software: MP, DM, Supervision: GA, LD, AG, TD, LWJB, CIDZ, Validation: AA, JK, AJM, LEAM, ZEK, SG, Visualization: NES, JEVB, JOA, AJM, Writing – Original Draft Preparation: NES, JEVB, GA, LD, Writing – Review & Editing: NES, JEVB, JOA, AJM, LB, VR, MF, DM, AG, TD, LWJB, CIDZ, LD, GA

## Acknowledgements

We thank Gabriele Köpf, Diana Giesler, Simon Köpf, Anke Biczysko and Beata Lukaszewska-McGreal for their excellent technical assistance, as well as the animal caretakers at Goethe University Frankfurt. Oli-neu cells were kindly provided by Prof. Jacqueline Trotter.

## Figures

**Supplementary Fig. S1:**
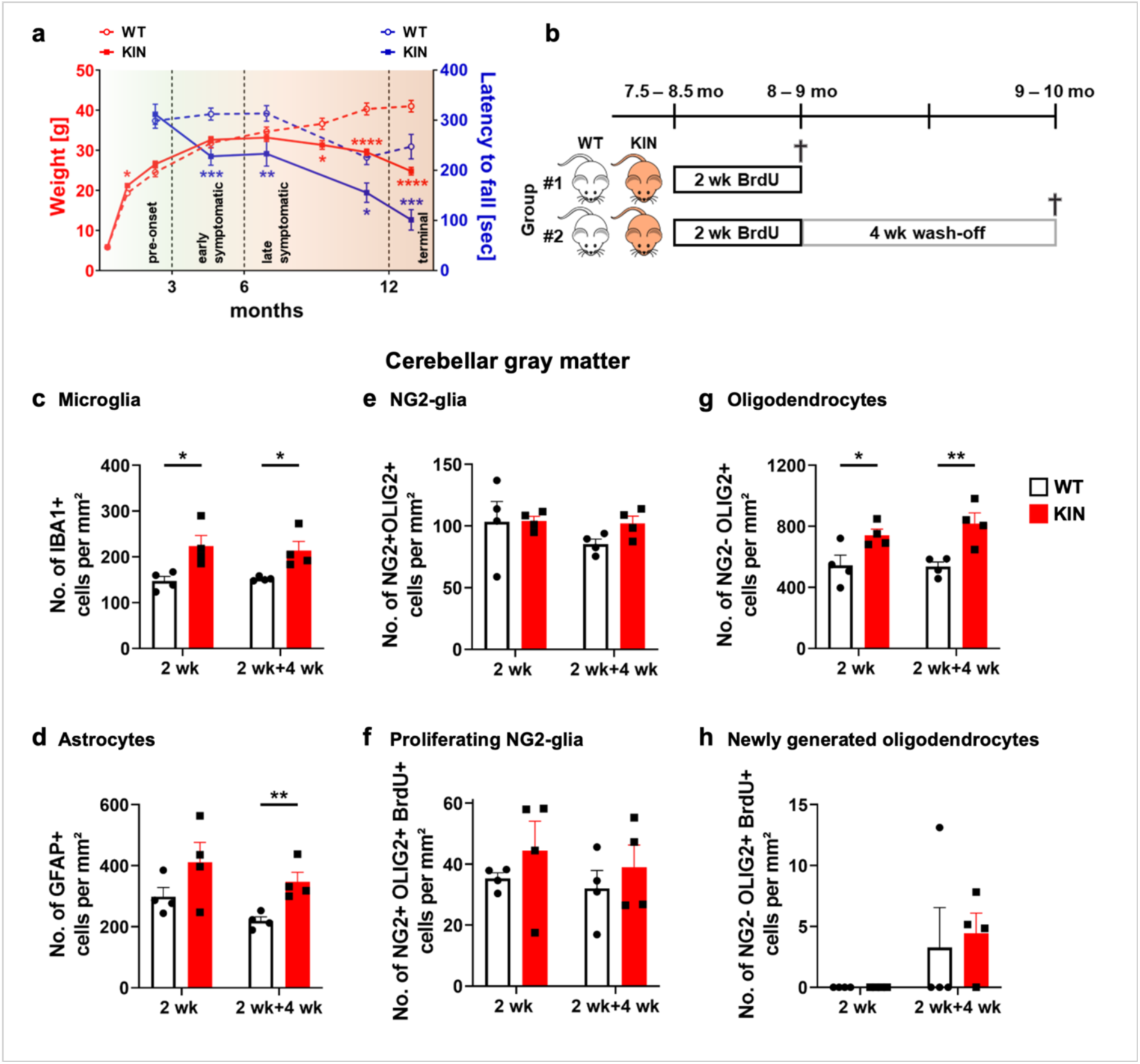
Analysis of microglial activation, astrogliosis and oligodendrocyte renewal in cerebellar gray matter. **a,** Body weight and Rotarod performance profiles (mean ± sem) of WT and KIN animals were depicted across lifespan in groups of 8-34 mice. Raw data were obtained from Sen *et al.*^18^ and graphed again covering both genders and phenotypic features to clarify the disease course. **b,** Scheme depicting the course of bromodeoxyuridine (BrdU) treatment of WT and KIN animals in late symptomatic stage. First experimental group was sacrificed immediately after the 2-week BrdU treatment supplied in drinking water, while the second group was sacrificed following an additional 4-week wash-off period to trace the progeny of labelled cells. Immunohistochemical analyses of cerebellar GM revealed significantly increased microglial activation **(c)** and astrogliosis **(d)** in KIN tissue in both treatment groups. While the number of total or proliferating NG2-glia were comparable **(e, f)**, KIN animals showed significantly higher number of mature oligodendrocytes in both BrdU treatment groups **(g)**. KIN animals also showed an overall increase of newly generated oligodendrocytes despite the high variation **(h)**. Each data point represents a single animal.

**Supplementary Fig. S2:**
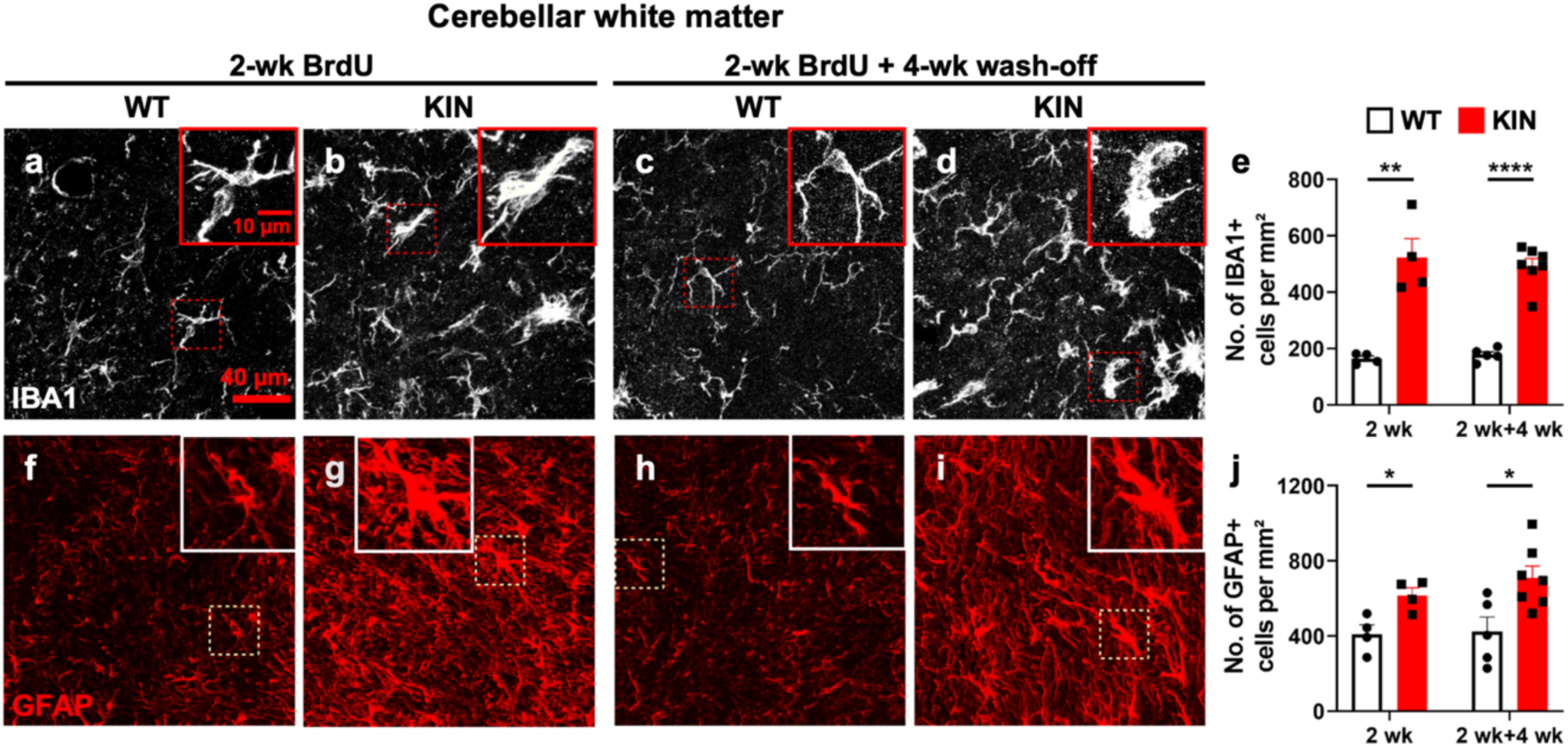
Increased microglial activation and astrogliosis in KIN cerebellar white matter. Immunohistochemical analyses of cerebellar white matter revealed significantly increased microglial activation upon IBA1 quantification (**a-e**) and astrogliosis upon GFAP quantification (**f-j**) in KIN tissue in both experimental groups treated with BrdU. Each data point represents a single animal.

**Supplementary Fig. S3:**
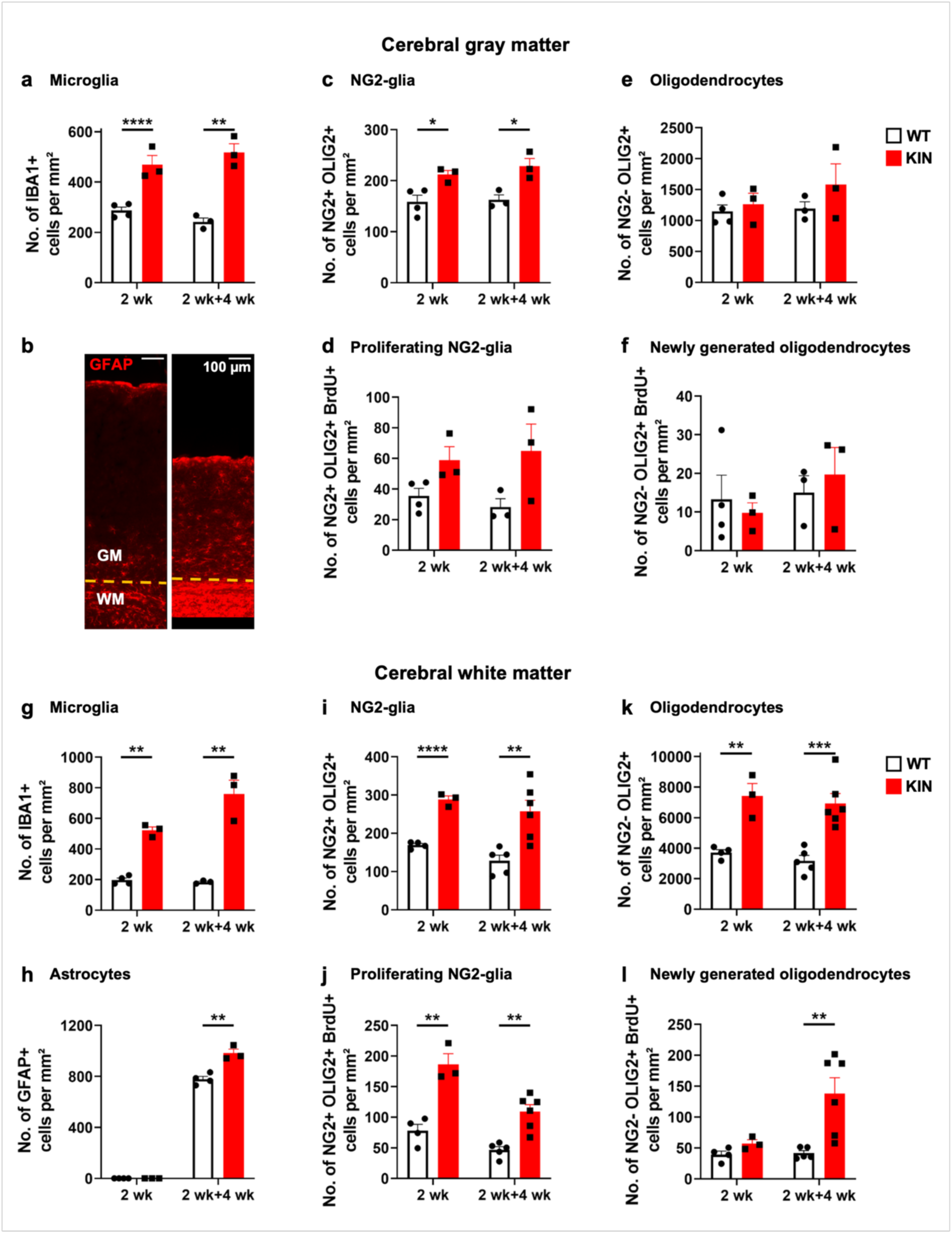
Quantification of microglial activation, astrogliosis and oligodendrocyte renewal in cerebrum. Immunohistochemical analysis and quantification of the cerebral GM **(a-f)** and WM **(g-l)** of WT and KIN mice show increased microgliosis **(a,g),** astrogliosis **(b,h)**, and accumulation of NG2- glia **(c,d,i,j)** in KIN tissue in both experimental groups treated with BrdU. Increased number of oligodendrocytes were only observed in the WM of KIN brain slices **(e,k)** and enhanced differentiation was only observed in the WM of KIN animals after 2wk+4wk BrdU treatment **(f,l)**. **b,** Representative tile-scan image of the cortical GM and WM, immunolabeled against GFAP (red). Each data point represents a single animal.

**Supplementary Fig. S4:**
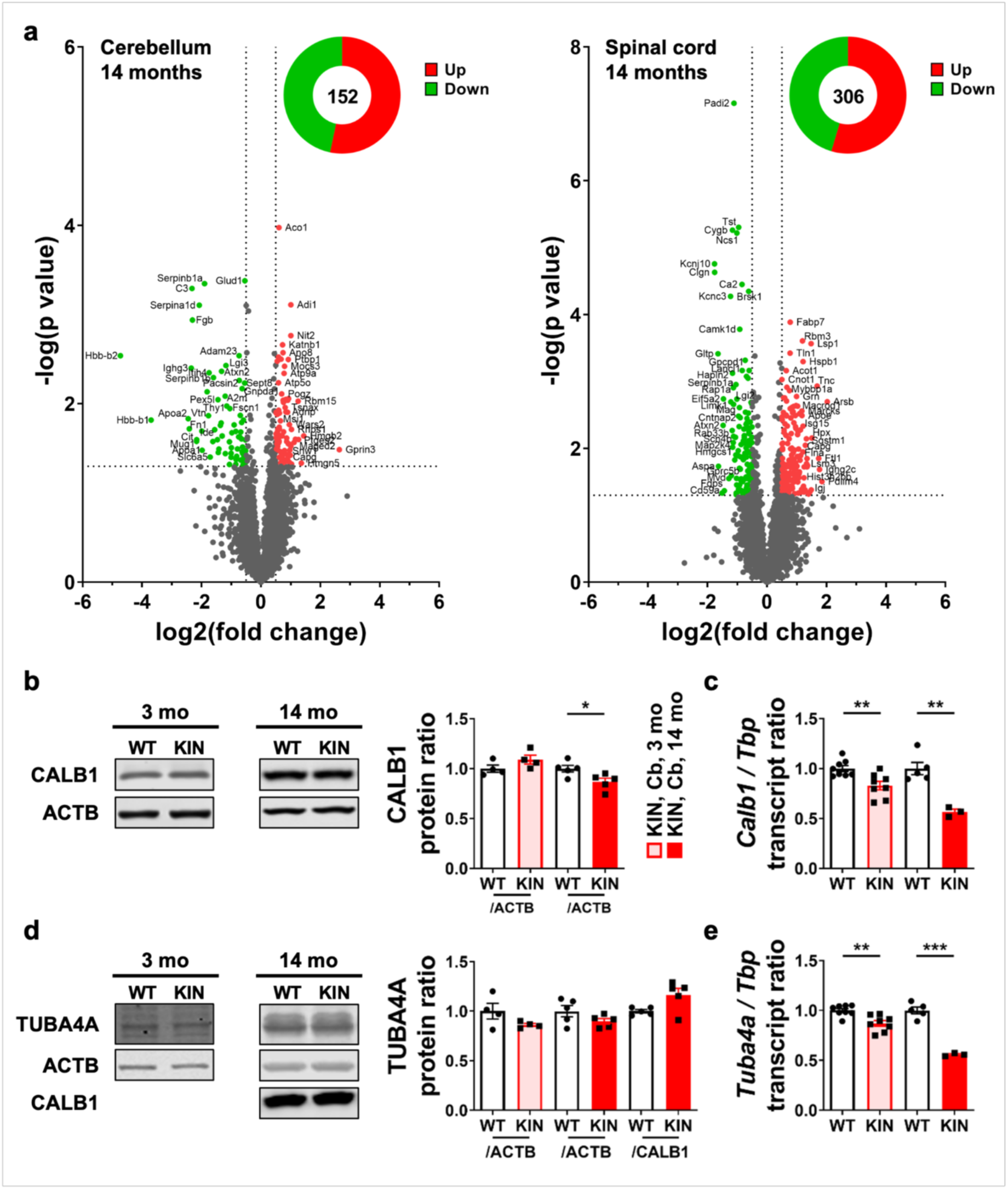
Cerebellum and spinal cord proteome profiles, and assessment of neuronal somatodendritic integrity. **a,** Volcano plots depict the label-free proteome data obtained from cerebellum and spinal cord tissues from WT and KIN animals at terminal stage (14 mo), identifying 3593 and 3769 proteins, respectively. Significantly down- and upregulated proteins in each dataset were labeled and colored in green and red, respectively. Protein levels of Calbindin-1 (CALB1) **(b)** as a marker of intact Purkinje neuron soma and TUBA4A **(d)** as a marker of intact somatodendritic compartment were measured by quantitative immunoblots in WT and KIN cerebellum (Cb) at pre-onset (3 mo) and terminal (14 mo) stages, showing only a mild downregulation of CALB1 at the terminal stage in KIN tissue. ACTB was used as a general loading control, and CALB1 was used as a normalizer to examine dendritic affection versus cell soma. Transcript levels of *Calb1* **(c)** and *Tuba4a* **(e)** were measured by qRT-PCR in WT and KIN cerebellum at pre-onset (3 mo) and terminal (14 mo) stages, showing a mild yet significant downregulation at pre-onset stage that progressively decreases at terminal stage in KIN tissue for both transcripts. *Tbp* was used as housekeeping gene. Each data point represents a single animal.

**Supplementary Fig. S5:**
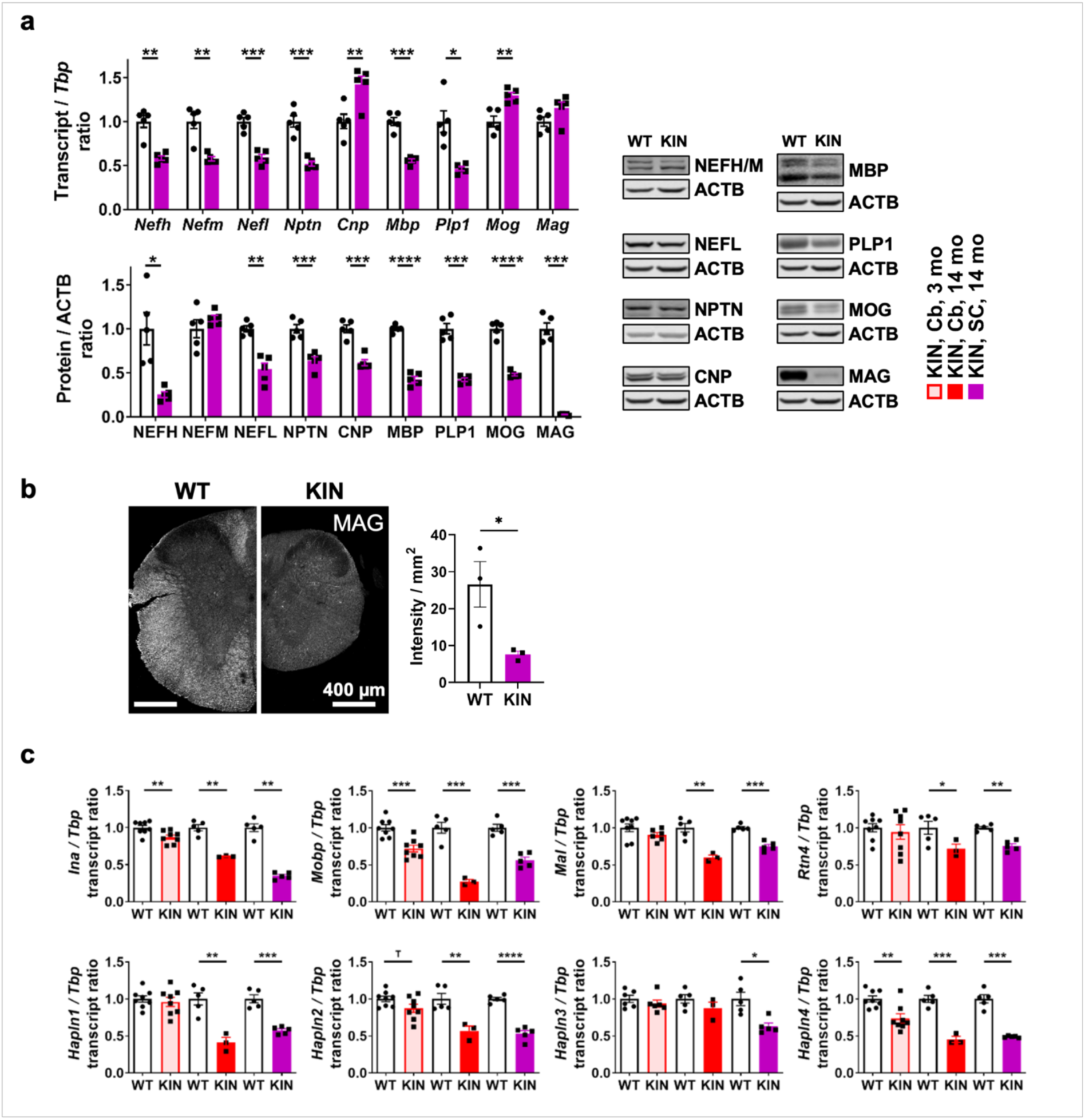
Assessment of demyelination in CAG100-KIN spinal cord with additional myelin markers. **A,** Transcript and protein levels of neuronal and myelin compartment factors in WT and KIN spinal cord at the terminal stage (14 mo), showing a dysregulation pattern similar to that of cerebellum. Reduced abundance of many neuronal and myelin proteins is mirrored by their transcript level, except the prominent downregulations of CNP, MOG and MAG proteins without a reduction of their transcripts. ACTB was used as loading control in quantitative immunoblots. *Tbp* was used as housekeeping gene in qRT-PCR experiments. Each data point represents a single animal. **b,** Immunohistochemical assessment of MAG protein in spinal cord sections from WT and KIN animals at terminal stage showed a marked reduction in KIN in agreement with immunoblot results. Each data point represents a technical replicate. **c,** Transcript levels of additional neuronal and myelin compartment factors in WT and KIN cerebellum (Cb, 3 mo and 14 mo) and spinal cord (SC, 14 mo), display early and progressive dysregulations of neuronal *Ina*, oligodendroglial *Mobp* and neuron-secreted *Hapln4*. Later onset dysregulations were observed in oligodendroglial *Mal*, *Rtn4*, and neuron-secreted *Hapln1-3*. *Tbp* was used as housekeeping gene in qRT- PCR experiments. Each data point represents a single animal.

**Supplementary Fig. S6:**
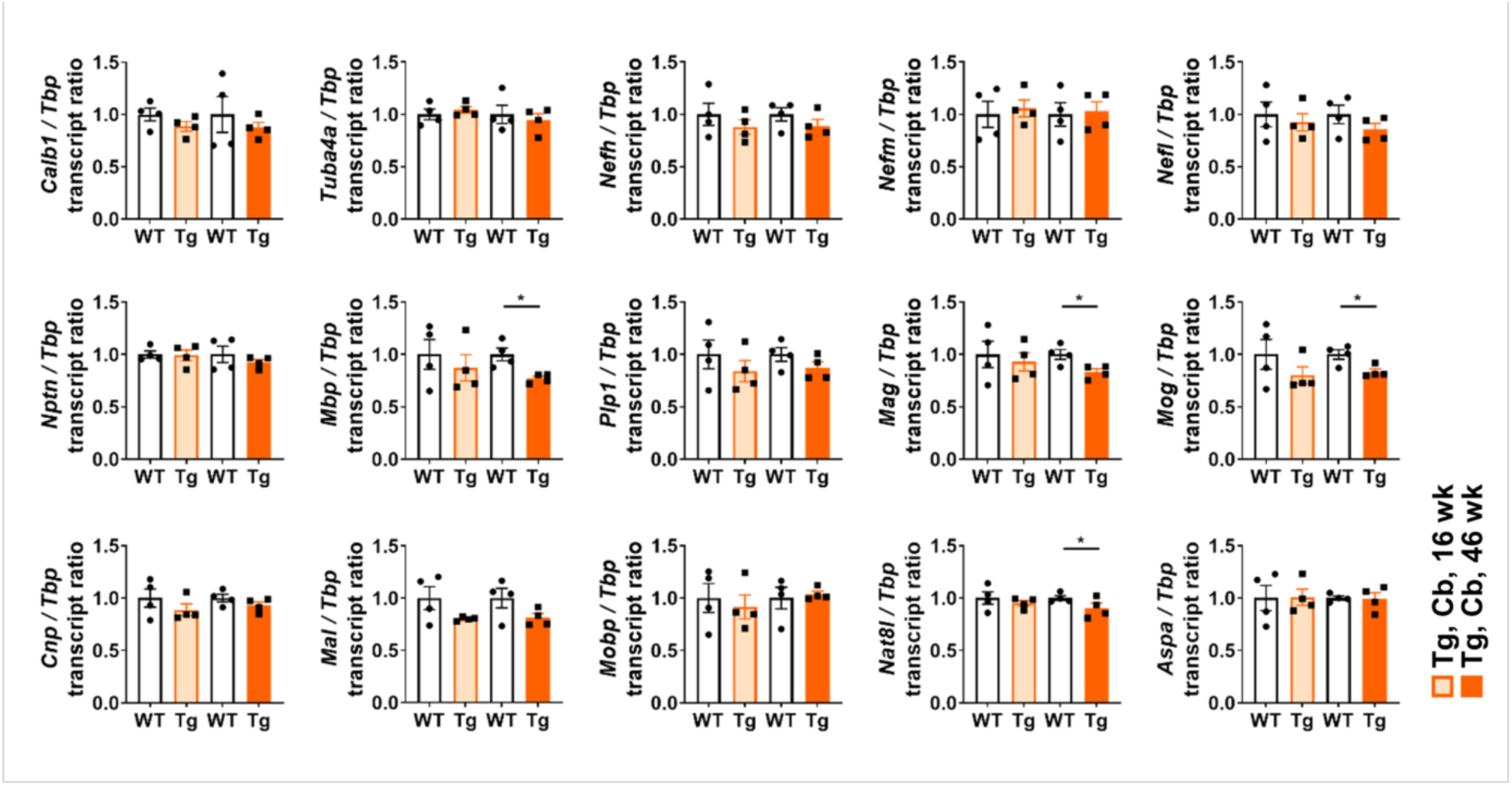
Assessment of demyelination *ATXN2*-Q58 transgenic mouse. Transcript levels of all neuronal and myelin compartment factors in WT and transgenic (Tg) *ATXN2*-Q58 mouse cerebellum at pre-onset (16 wk) and late-symptomatic (46 wk) stages. While no dysregulation of axonal markers was observed in early or late disease stages, only a mild downregulation was observed in several oligodendroglial transcripts, such as *Mbp*, *Mag*, *Mog* and *Aspa.* The dysregulation pattern of *Mag* and *Mog* were found opposite to that of KIN mice at this stage. *Tbp* was used as housekeeping gene in qRT-PCR experiments. Each data point represents a single animal.

**Supplementary Fig. S7:**
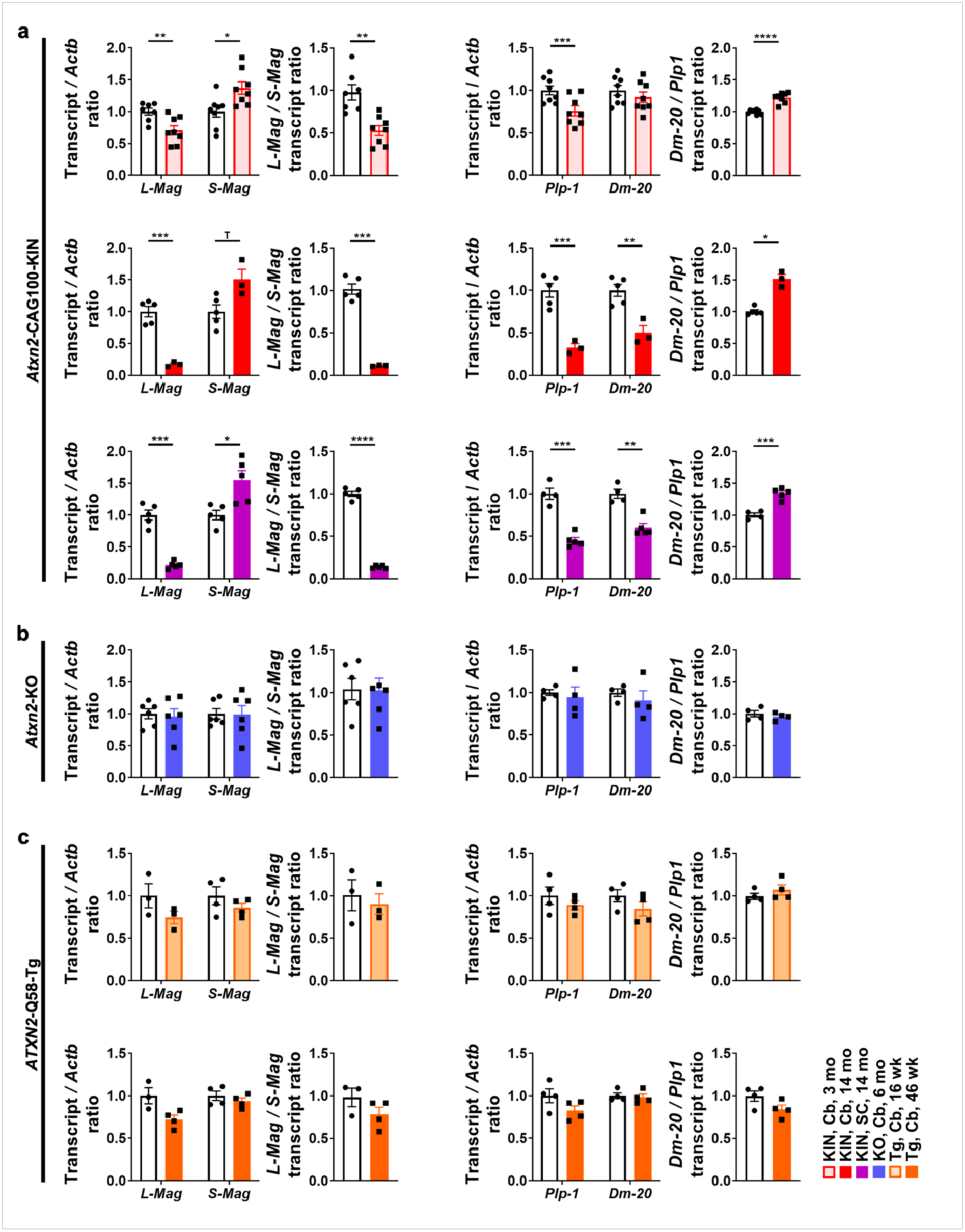
Validation of *Mag* and *Plp1* splicing defects with an independent technique confirms the impact of ATXN2 toxicity in oligodendrocytes. Transcript levels of *Mag* and *Plp1* splice isoforms were measured by qRT-PCR in **(a)** KIN cerebellum (Cb, 3 mo, 14 mo) and spinal cord (SC, 14 mo), **(b)** KO cerebellum (Cb, 6 mo) and **(c)** transgenic (Tg) *ATXN2*-Q58 mouse cerebellum (Cb, 16 wk, 46 wk) in comparison to age-matched WT controls. The splicing defects previously observed with semi-qPCRs (Figure 6) were validated for all KIN tissues. No splice defects were observed in KO or Tg *ATXN2*-Q58 cerebellum, suggesting that ATXN2 aggregation toxicity specifically in oligodendroglia impacts the splicing pattern of mature myelin markers. *Actb* was used as housekeeping gene in qRT-PCR experiments. Each data point represents a single animal.

**Supplementary Fig. S8:**
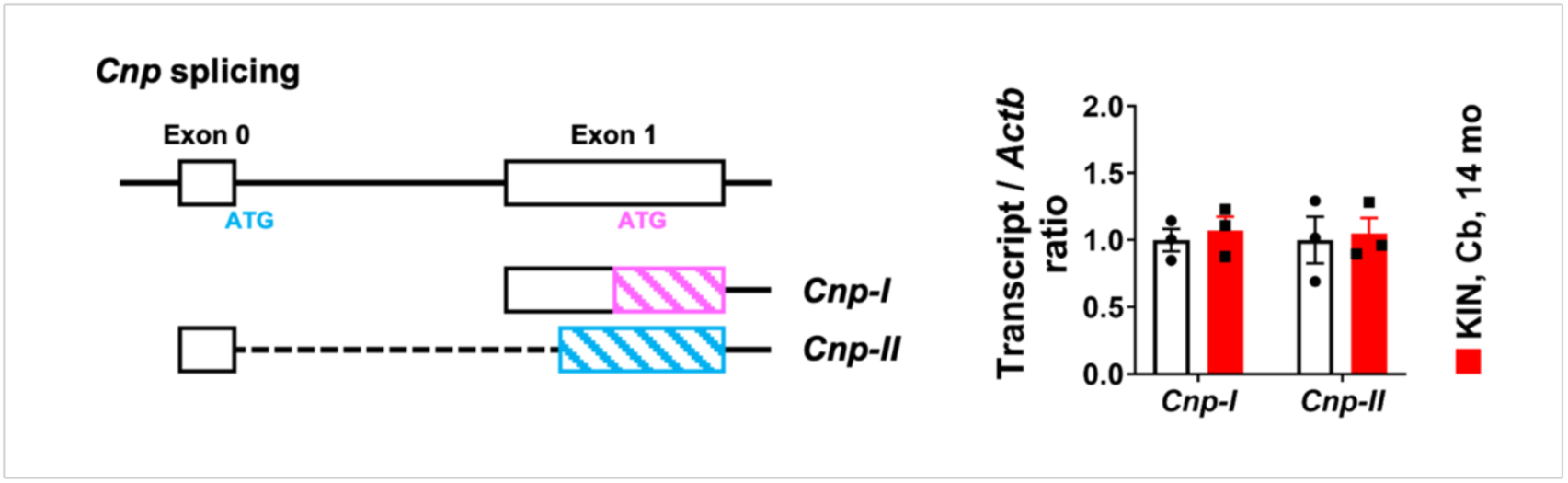
Assessment of *Cnp* splicing. Expression levels of *Cnp* transcript isoforms, produced by an unknown splice factor, were measured by qRT-PCR in KIN cerebellum (Cb) at 14 months. No alteration of its splicing pattern was observed even at the terminal disease stage, suggesting that the alterations of *Mag* and *Plp1* levels observed earlier were specific outcomes of ATXN2 pathology. *Actb* was used as housekeeping gene. Each data point represents a single animal.

**Supplementary Fig. S9:**
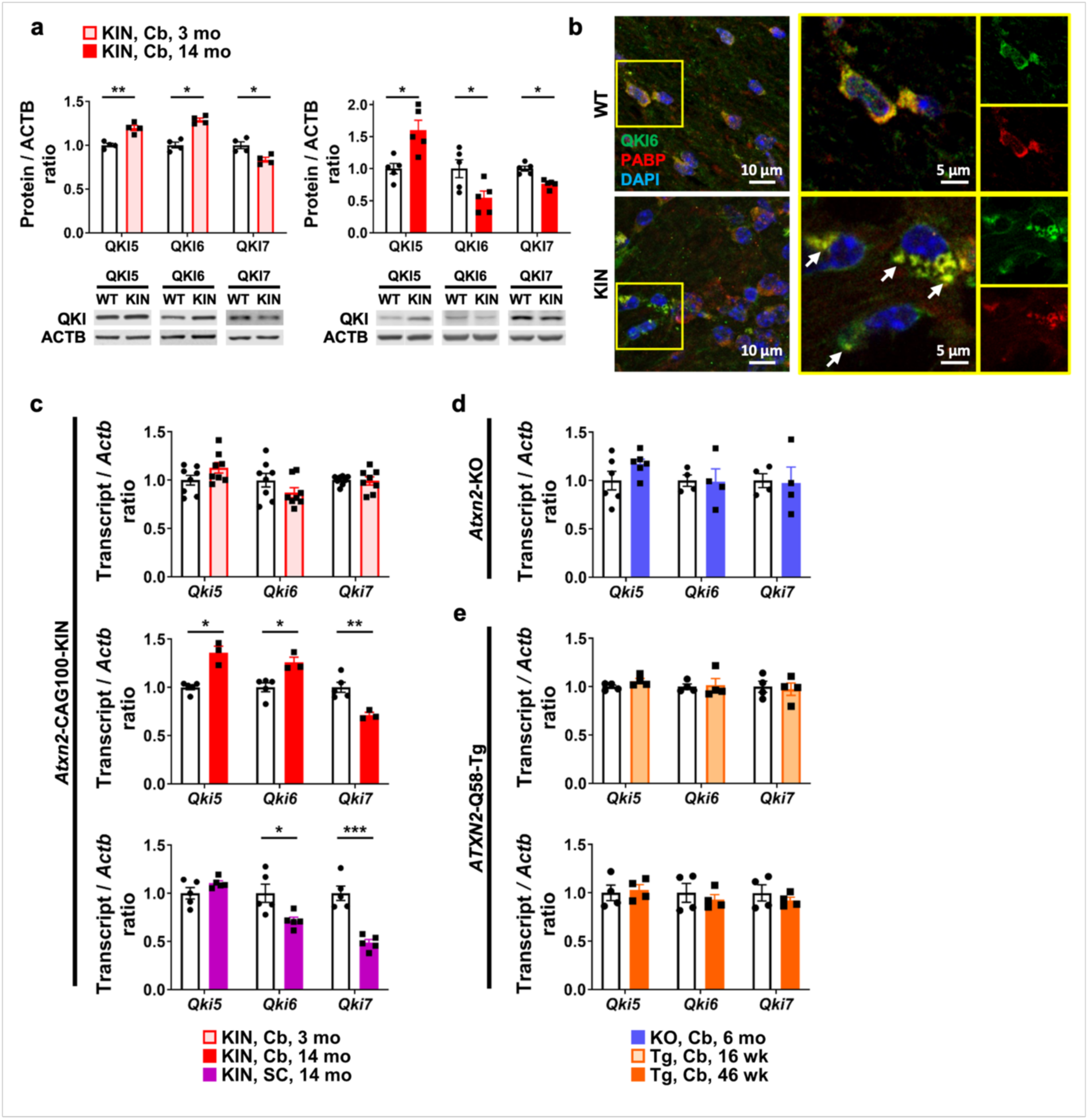
Protein and transcript levels of *Qki* splice isoforms are differentially affected by ATXN2 pathology. **a,** Protein levels of QKI5, QKI6 and QKI7 were measured in KIN cerebellum (Cb) at pre-onset (3 mo) and terminal (14 mo) disease stages with age-matched WT controls. QKI5 remained upregulated and QKI7 remained downregulated in KIN tissue throughout the disease course, but QKI6 showed a pre-onset upregulation that transformed into a strong downregulation at the terminal stage, mimicking a profile usually displayed by proteins sequestrated into aggregation in the disease course. ACTB was used as loading control in quantitative immunoblots. Each data point represents a single animal. **b,** Immunohistochemical assessment of QKI6 (green) localization in WT and KIN cerebellar WM at the terminal stage (14 mo) showed its diffuse cytosolic distribution in WT samples, and its sequestration into cytosolic aggregates in KIN samples marked by SG marker PABP (red). DAPI (blue) marks the nuclei. Transcript levels of *Qki5*, *Qki6* and *Qki7* were measured with specific primers designed for qRT-PCR in KIN **(c)**, KO **(d)** and transgenic (Tg) *ATXN2*-Q58 mouse **(e)** tissues at different disease stages with age-matched WT controls. Only late-onset dysregulations were observed in KIN cerebellum (Cb) and spinal cord (SC), without a change in KO or Tg *ATXN2*-Q58 cerebellum. *Actb* was used as housekeeping gene in qRT-PCR experiments. Each data point represents a single animal.

